# A basic leucine zipper uses a dimer pathway to locate its targets in DNA mixtures

**DOI:** 10.1101/2025.08.25.672131

**Authors:** Mikhail Kuravsky, Gabriel Grey, Conor Kelly, Polenus Sarkar, Jeremie Gaudez, Sarah L. Shammas

## Abstract

The operation of Eukaryotic transcription factors remains enigmatic. Cyclic AMP-responsive element-binding protein (CREB) is a member of the basic zipper family, a superfamily of transcription factors which operate exclusively in eukaryotes and bind DNA targets as homodimers or heterodimers. Modulation of oligomerization provides an additional opportunity for transcriptional control by this (and similar) families over monomeric transcription factors. However, when dimerization occurs – before or after target binding – is not known. We performed a suite of *in vitro* stopped-flow kinetic measurements, including CREB basic zippers target search amongst excess non-target DNA. The extensive dataset enabled a kinetic and thermodynamic understanding of DNA binding that demonstrated most productive search is performed by dimeric, rather than monomeric, CREB. Equilibrium is approached very rapidly under physiologically relevant concentrations, where relative flux through the monomer pathway is only around 1 in every 10,000 complexes formed. This preference of mechanism is driven by CREB monomer having a substantially higher affinity for another CREB monomer than for its DNA target. Equilibrium experiments with eight other monomeric peptides further suggest this as a common feature amongst the bZIP proteins, with only one peptide (Jun) displaying similar affinities for both. The work has implications for understanding the nature of DNA target search, as well as designing efficient artificial transcription factors and transcriptional inhibitors.

## INTRODUCTION

Within the nucleus, specific transcription factors find their cognate binding sites amongst huge excesses of non-target DNA. Target search can be complex and involves several processes, including 3-dimensional diffusion, DNA sliding, hopping and jumping, and intersegmental transfer^1–3^.

Nonetheless much progress has been made in understanding these phenomenon, especially within bacterial systems^4^. This experimental work has been performed with monomeric transcription factors or preformed (stable) oligomeric transcription factors^5^, which is understandable given the search process is already hugely complex, and suitable in that the majority of transcription factors bind their targets as monomers. However many eukaryotic transcription factors are oligomeric^6,7^, notable examples being the trimeric heat shock protein transcription factors^8^ and the dimeric basic helix-loop-helix (bHLH) and basic zipper (bZIP) superfamilies^9,10^. Roughly 10% of human transcription factors contain one of the latter two domains^7^.

Co-operativity of binding between various members of transcriptional complexes is thought to be important for accomplishing the specificity required for successful transcriptional regulation^7,11^. Interestingly artificial introduction of oligomerisation domains into transcription factors can enhance specificity and sensitivity, including within mammalian cells^12,13^. Theoretical work suggests oligomerisation of DNA-binding proteins can enhance or impair both target-site occupancy and dwell times, as well as redistribute the proteins spatially on DNA^14^. It has been further suggested that the kinetic stability of protein oligomers on DNA may have an optimal value for efficient target search, where both monomers and dimers are present and sliding on DNA^15^. Modulation of oligomeric state within the nucleus, for example by phosphorylation^16^ presents an opportunity to control transcriptional activity. These observations highlight the importance of identifying the order of oligomerization and target binding in understanding the more complex search process of oligomeric transcription factors. An oligomeric transcription factor, signal transducer and activator of transcription 3 (STAT3), has been shown to display oligomeric state-dependent mobility within the nucleus^17^. STAT3 was shown to enter the nucleus as dimers, and then form tetramers upon binding to DNA in a fashion that depends upon chromatin accessibility. Assessing stoichiometry within cells can already be a technical challenge, so uncovering the stoichiometry as it binds to a target is even more so. Single-molecule tracking inside cells and in a reconstituted in vitro system was used to propose that the bHLH pioneer transcription factor Sox2 binds to DNA and slides as a monomer, with Oct4 heterodimerization occurring after it has found its target^18^. However, subsequent analysis of their data by Biddle *et al*. argues that the order of binding events was not revealed, but instead the work uncovered negative reciprocity of DNA binding of the two factors^19^, plausibly through changes in chromatin accessibility^20^.

The basic leucine zipper (bZIP) family is a dimeric eukaryotic superfamily of transcription factors and includes 56 human members^21,22^. Structures have been solved for many bZIP proteins that show them bound to their DNA targets as coiled-coil dimers, with their helices inserted into DNA major grooves like a pair of chopsticks (Fig. 1); DNA binding is mediated by a basic region (BR), with a leucine zipper (LZ) mediating dimerization^10,23^. One prominent member of the family is the cAMP-response element binding protein (CREB). Originally identified as a TF that activates expression of the somatostatin gene in response to cAMP signalling^24^, it was subsequently discovered to regulate a much wider range of functions, playing a pivotal role in controlling proliferation^25–27^, differentiation^28– 30^ and survival of cells^31,32^, and memory formation^33–35^. The binding of CREB to its target cAMP-response element sites (consensus CRE, 5’-TGACGTCA-3’) is through its C-terminal basic leucine zipper (bZIP) domain^36^.

**Fig. 1.**
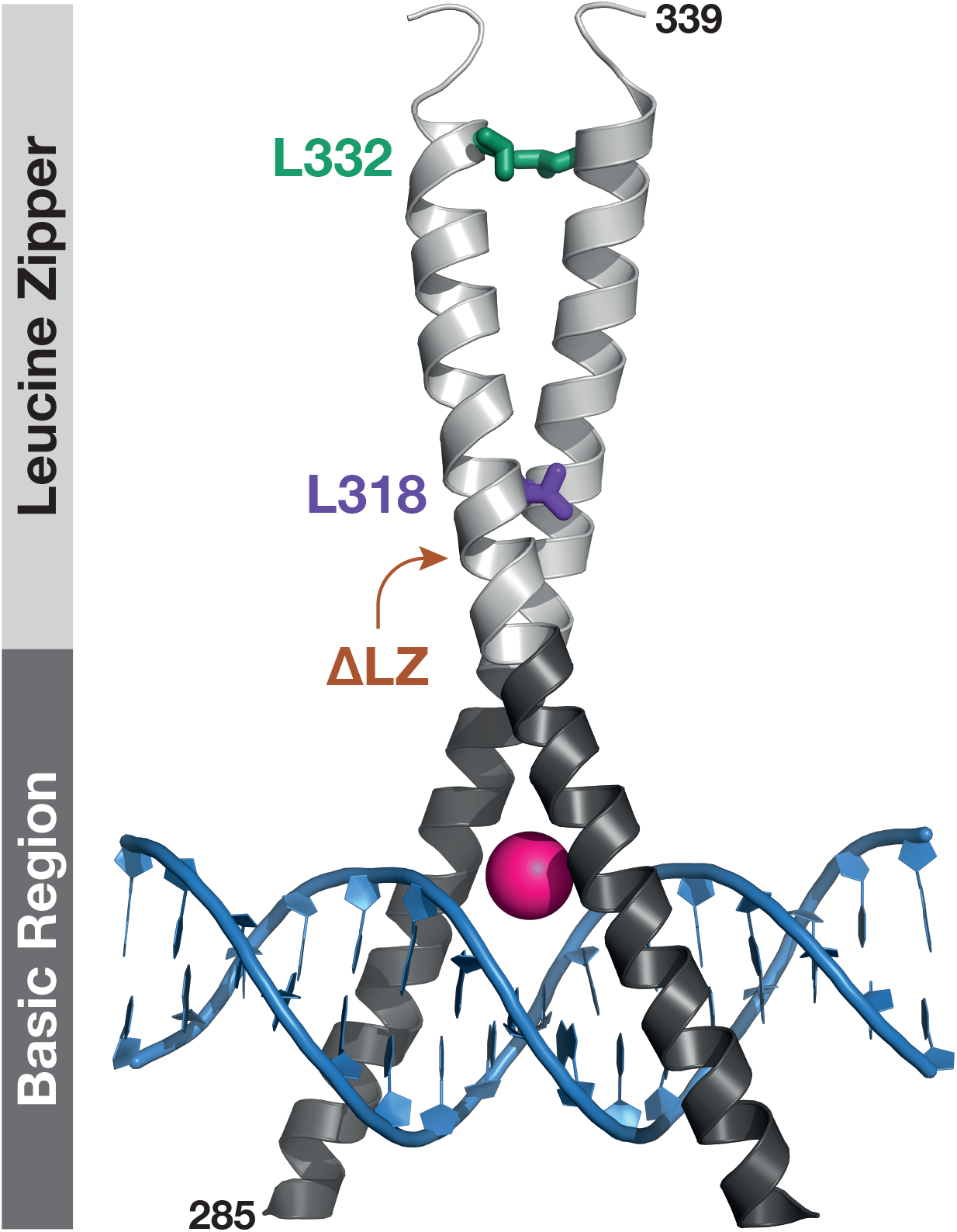
Structure of the complex between CREB bZIP (grey) and CRE DNA (blue). CREB adopts a typical bZIP conformation, inserting α-helices within the major grooves of the DNA. Positions of leucine residues subjected to mutagenesis to destabilise the dimer are shown as coloured sticks. The truncation position for generating the monomeric CREB is also shown (brown). Pink sphere represents a magnesium ion. The figure is derived from PDB 1DH3^23^. Note that residues 277-285 are not shown because they were not present in the X-ray crystallography construct.

Dimerization preferences of the human bZIPs have been well researched, most notably in pioneering studies by the Keating lab^22,37,38^, and the “rules” governing the preferences are understood sufficiently well for predictions and designs to be made^37,39,40^. However the mechanisms by which the bZIP proteins find their DNA targets are less well elucidated, and importantly different bZIPs may not necessarily follow the same pathway. Limiting cases where proteins first bound DNA and then dimerised (monomer pathway) or dimerised and then bound DNA (dimer pathway) were considered (Fig. 2a). Attempts have been made *in vitro* to identify the stoichiometry of a limited number of bZIP domains in the “search state” and studies proposed a monomer pathway during search^41–48^. Firstly, a pre-requisite for the monomer pathway is the existence of DNA bound monomers, and these have been directly observed for some bZIPs. For example, the basic region of plant AtbZIP16 binds DNA in EMSA assays^49^ and the basic region of v-Jun protected its DNA target site in footprinting assays^47^. Secondly, the monomer pathway is considered favoured kinetically. I*n vitro* kinetic studies conducted using the yeast bZIP GCN4 showed that rate constants for DNA binding are orders of magnitude higher than those for dimerization^41^. Heterodimerization of its mammalian homolog Jun with its partner bZIP Fos is accelerated by DNA, with monomer pathways having a statistically superior description of the data^44^.

**Fig. 2.**
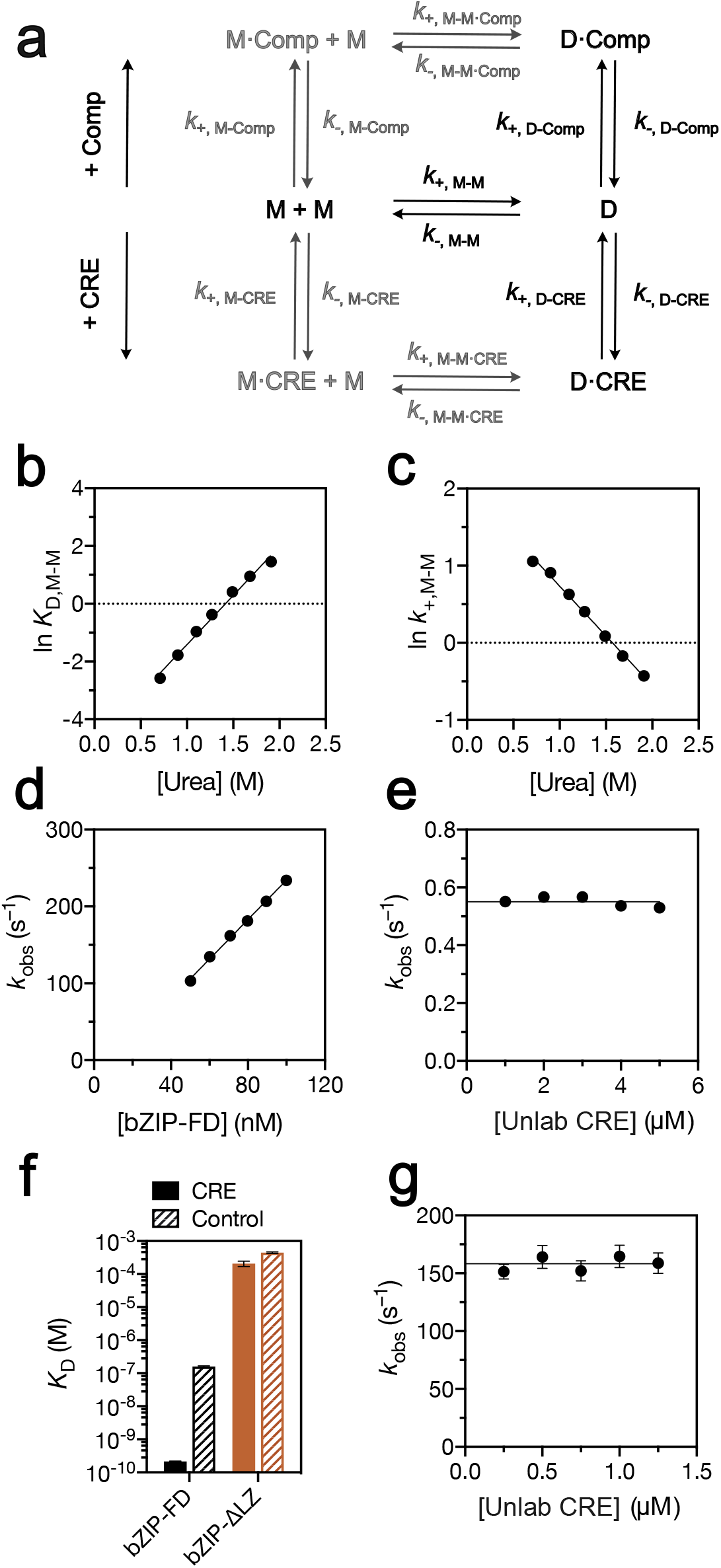
Development of a simple kinetic scheme for target search. **a**, Simplified reaction scheme describing the formation of complexes between CREB bZIP, target DNA (CRE) and competing non-target DNA (Comp). The rate constants are named according to the two molecules (separated by −) which are associating (*k*_+_) or separating from each other (*k*_−_). Molecules joined by · remain bound throughout the elementary reaction. Equilibrium constants in the text use the same terminology for consistency. M, monomeric CREB bZIP; D, dimeric CREB bZIP; *CRE, CRE* DNA; Comp, non-target DNA. Reactions modelled within the dimer-only pathway are shown in black, and those neglected in a dimer-only binding pathway are grey **b, c**, Equilibrium dissociation constants for monomer-dimer transition *K*_D, M-M_ **(b)** and association rate constants for protein dimerization *k*_+,M-M_ **(c)** determined from stopped-flow fluorescence kinetic refolding experiments performed in the presence of varying concentration of urea. The lines are the best fits to a linear function; extrapolation of fits to the zero urea allows estimation of *K*_D, M-M_ and *k*_+,M-M_ in the absence of denaturant. **d**, *k*_obs_ values for the association of CREB-bZIP-FD with Alexa Fluor® 488-CRE measured under the pseudo-first order conditions with an excess of protein. Gradient of the linear fit corresponds to the association rate constant *k*_+,D-CRE_. **e**, *k*_obs_ values for the dissociation of CREB-bZIP-FD and Alexa Fluor® 488-CRE measured in out-competition experiments, in which unlabelled CRE was used to displace the Alexa Fluor® 488-CRE from the complex. The dissociation rate constant *k*_−,D-CRE_ is estimated by averaging *k*_obs_ values. **f**, Equilibrium dissociation constants with Alexa Fluor® 488-CRE and Alexa Fluor® 488-Comp DNA determined for the disulphide-linked CREB-bZIP-FD (*K*_d, D-CRE_ and *K*_d, D-Comp_) and a monomeric version without the leucine zipper CREB-bZIP-ΔLZ (*K*_d, M-CRE_ and *K*_d, M-Comp_). Values determined from equilibrium binding titrations, except for *K*_d, D-CRE_ which was estimated as a ratio of *k*_−,D-CRE_ to *k*_+,D-CRE_. **g**, *k*_obs_ values for the dissociation of the CREB-bZIP-FD· Alexa Fluor® 488-Comp complex measured in out-competition fluorescence anisotropy stopped-flow experiments. *k*_−,D-Comp_ was calculated as an average of individual *k*_obs_ values. All errors are the standard errors of fit.

Although existence of monomeric intermediates is necessary, it is not sufficient for identifying the active search species as monomer. Whilst folded as a dimer bound to DNA, bZIP domains are understood to be disordered in isolation^10^. The original pathway studies^41–46^ were performed with a limited number of related bZIP domains and largely concentrated on whether the protein was folding and dimerising by itself, or on DNA. Several studies monitored dimerization, and their results were consistent with dimerization being accelerated by the presence of DNA (and other polyanions)^43^. This is distinct from describing how target search occurs in DNA mixtures since dimer might be formed on target sequences (monomer pathway) or on non-target DNA and then at some time later bind the DNA target (dimer pathway). Nonetheless reference to the kinetic favouring of a monomer pathway in target search continue, in some cases despite acknowledgement of high entropic costs for specific monomer-DNA interactions that arise from a requirement to fold upon binding^5,10,49–68^. Our recent kinetic studies with a CREB bZIP construct demonstrated that CRE binding by CREB dimer is rapid and electrostatically enhanced, with a relatively early transition state^69^. CREB bZIP itself has not been the focus of any kinetic pathway studies, but it has until recently been considered even more likely to pursue a monomeric search mode due to reports of very high equilibrium dissociation constants, in the tens of micromolar, for homodimerization^70^. Recently, publications from ourselves^69^ and another group^71^ have found much higher binding affinities.

Here we present kinetic and equilibrium studies of DNA target binding by CREB, including ones performed in the presence of large excesses of non-target DNA, that were designed to reveal the stoichiometry of the search pathway for CREB. Our results challenge the existing narrative of a monomer search pathway, and importantly are able to rationalise the preferred dimeric pathway in terms of DNA binding thermodynamics and specificity. Appreciation of the underlying principles determining preferred search stoichiometry may assist design of artificial transcription factors and transcriptional modulators, which might be valuable as therapeutics or as tools for functional studies in cells.

## METHODS

### Cloning and mutagenesis

Amino acid numbering of CREB in this paper is based on the isoform CREB-A. Plasmid encoding extended CREB-bZIP construct (residues 277-339) was produced from the _285_bZIP plasmid generated in our previous publication^69^. This plasmid contains chemically synthesized coding sequence for residues 285-339 bZIP domain of human CREB1 linked with the GB1 solubility-enhancement tag and inserted into a pRSET A vector containing a 6xHis tag at the N-terminus (Invitrogen). All cysteine residues in the bZIP domain (C300, C310 and C337) were replaced with serine residues. These mutations were previously reported to significantly improve the solubility of the protein without altering its structure and the affinity to the cognate DNA sequence^72^. A tobacco etch virus (TEV) protease site was placed between the GB1 tag and the bZIP sequence. The substitutions were introduced by Q5 site-directed mutagenesis (New England Biolabs). A C-terminal GGC extension was added to permanently join subunits of a dimer by an intermolecular disulphide bond (CREB-bZIP-FD). The L318V and L332V mutations aiming to destabilize the coiled-coil structure of the leucine zipper were produced from the CREB-bZIP plasmid without the GGC extension. Plasmid encoding a shortened version of CREB-bZIP lacking the most part of the leucine zipper (CREB-bZIP-ΔLZ, residues 277-315) was obtained by introducing a nonsense A316* substitution. To generate plasmids encoding basic-region only peptides of other human bZIP domains, the bZIP 277-339 sequence was replaced using Gibson assembly. The 277-339 sequence was removed by Polymerase Chain Reaction using primers (Life technologies) and Q5 hot-start polymerase utilizing manufacturer’s instructions (New England Biolabs (NEB)) to generate the backbone template. DNA sequences corresponding to forced monomer sequences with flanking sequences were purchased as gBlocks Gene Fragments (Integrated DNA Technologies), and inserted with NEBuilder® HiFi assembly mix (New England Biolabs (NEB)) as per manufacturer instructions. All plasmids were verified by DNA sequencing.

### Protein expression and purification

Plasmids encoding the protein constructs were transformed into *Escherichia coli* BL21(DE3)pLysS. The cells were cultured in 2xYT medium supplemented with 0.2% (w/v) glucose until the optical density (OD_600_) reached 0.6-0.8. Protein expression was induced with 1 mM IPTG, and the culture was grown for further 4 h at 37 ºC. Cells were harvested by centrifugation and lysed by sonication in buffer A (100 mM Tris-HCl pH 8.5, 50 mM NaCl). The lysates were cleared by centrifugation and loaded on HisTrap HP columns (GE Healthcare) equilibrated in buffer A. The columns were consecutively washed with buffer A and buffer B (100 mM Tris-HCl pH 8.5, 2 M NaCl), and the proteins were eluted with a 0-1.5 M gradient of imidazole in buffer A. The fractions containing 6xHis-GB1-bZIP were pooled and exchanged into buffer A supplied with 1 mM EDTA and 1 mM DTT in the presence of 2 µM TEV protease. Due to the TEV cleavage an N-terminal glycine residue is present in all purified peptides. To separate the cleaved 6xHis-GB1 tag, the protein was loaded on a HiTrap SP cation exchange column (GE Healthcare) equilibrated with buffer C (10 mM HEPES pH 7.5) and eluted with a 0-2 M gradient of NaCl in buffer C. To dimerise the constructs with the GGC extension, the proteins were concentrated to 4 mM or higher, mixed with an equal volume of 100 mM Tris-HCl buffer (pH 9.2), placed in the tubes with loose fitting caps and kept in a shaking incubator for 16 h at 37 ºC. The presence of disulphide bond was confirmed by SDS-PAGE and mass spectrometry (for example Fig. S2). The fractions containing pure bZIP were pooled, buffer exchanged using a 5 ml HiTrap desalting column into 10 mM MES pH 6.5, 150 mM NaCl, 10 mM MgCl_2_, 0.05% Tween-20, and then flash-frozen in liquid nitrogen and stored at −80 ºC until use. The purity of protein was verified by SDS-PAGE, and the absence of nucleic acid contamination was assessed by the ratio of absorbances at 260 and 280 nm.

### Design of forced monomer (basic-region only) sequences

In designing suitable sequences for forced monomer bZIP domains that lacked the leucine zipper, two considerations were made for extending the sequences further than the Uniprot defined basic region. Only human bZIPs with structural data were considered. Additional residues were added to the N-terminus until all contact points and stabilising structure observed in the crystal structure were included. Predicted helicity of the sequences was also considered since residual helical content might control DNA binding affinity through entropic requirements for folding upon binding. A series of sequences for each basic region was considered with differing (consecutively earlier) N-terminal starting residues. Helicity predictions were made for each were made using AGADIR algorithm^73^ (Centre of Genomic Regulation). Parameters for predictions were: pH = 7.4, temperature = 298 K, ionic strength = 0.15. The sequence selected was the shortest sequence which produced a predicted helicity similar to the full length, and maintained a consistent prediction when additional N-terminal residues are added. A similar strategy was used to identify a suitable C-terminal end for the sequence, however to balance requirement for behaving as a forced monomer, no ends were placed more than five amino acids after the six amino acid bZIP spacer sequence (one residue before the first leucine of the leucine zipper). Multiple sequence alignment was done manually, and with reference to crystal structures. Following identification of suitably truncated constructs all 54 known human bZIP domains could be successfully aligned using EMBL-EBI’s Clustal omega^74^.

### Forced monomer (bZIP-ΔLZ) protein expression and purification

Plasmids encoding the protein constructs were transformed into Escherichia coli BL21(DE3)pLysS. The cells were cultured in 2xYT medium supplemented with 0.2% (w/v) glucose until the optical density (OD600) reached 0.6-0.8. Protein expression was induced with 1 mM IPTG, and the culture was grown for further 4 h at 37 ºC. Cells were harvested by centrifugation with pellets being store at – 80 ºC until purification. The cells were lysed by sonication in buffer D (100 mM Tris-HCl pH 8.5, 50 mM NaCl). The lysates were cleared by centrifugation and loaded on HisTrap HP columns (GE Healthcare) equilibrated in buffer D. The columns were consecutively washed with buffer D and salt wash buffer (100 mM Tris-HCl pH 8.5, 2 M NaCl), and the proteins were eluted with a 0 – 500 mM gradient of imidazole in buffer D. The fractions containing 6xHis-GB1-bZIP-ΔLZ were pooled and exchanged into buffer E (10 mM sodium phosphate pH 8.0) supplied with 1 mM EDTA and 1 mM DTT in the presence of 10 µM TEV protease. Cleavage was overnight at room temperature. To separate the cleaved 6xHis-GB1 tag and any additional non-desired proteins, the protein was loaded onto a 5 mL HiTrap™ Heparin column (GE Healthcare) and then washed with 50 ml of 8M urea in buffer E. Protein was eluted with a 0-0.5 M gradient of sodium chloride in buffer E. Fractions were pooled and dialysed into buffer F (pH 7.0 10 mM HEPES) overnight at 4 ºC, and subsequently loaded onto a 5 mL HiTrap™ SP-HP column (GE Healthcare), washed with 50 ml of buffer F, and eluted with a shallow 0-2 M gradient of sodium chloride in buffer F. Fractions were analysed by SDS-PAGE and fractions without visible contaminants were pooled, buffer exchanged into 10 mM HEPES pH 7.4 150 mM NaCl 10 mM MgCl_2_ using a 5 ml HiTrap desalting column (GE Healthcare), flash-frozen in liquid nitrogen and stored at −80 ºC until use.

### Protein concentrations

The concentration of proteins was determined by absorbance measurements at 280 nm. The extinction coefficient for dimeric CREB-bZIP was estimated as 5120 ± 23 M^−1^cm^−1^ using the method described by Gill and von Hippel^75^. The extinction coefficient for dimerized constructs harbouring the GGC extension (CREB-bZIP-FD) was inferred to equal 5240 M^−1^cm^−1^, given the extinction coefficient of a free cystine (120 M^−1^cm^−1^)^76^. Extinction coefficients for monomeric constructs were calculated as: CREB - 1280 M^−1^cm^−1^, NFIL3 – 7442.04 M^−1^cm^−1^, CEBPA – 1280 M^−1^cm^−1^, CEBPB – 1166 M^−1^cm^−1^, JUN – 1282 M^−1^cm^−1^, JUND – 1400 M^−1^cm^−1^, ATF2 - 5114 M^−1^cm^−1^, MAFG – 1096 M^−1^cm^−1^, FOS - 1226 M^−1^cm^−1^. To achieve high accuracy in concentrations the samples were prepared gravimetrically for biophysical experiments.

### Preparation of double stranded DNA

To examine the binding of CREB to its cognate sequence we used a previously utilised self-annealing hairpin oligonucleotide^69,77,78^ (5’-CC<u>TGACGTCA</u>GCCCCC<u>TGACGTCA</u>GG-3’) labelled at the 5’-terminus with AlexaFluor® 488 (AlexaFluor® 488-CRE, Fig S2b). Dissociation experiments utilised an unlabelled double-stranded linear 14 bp oligonucleotide containing an 8 bp palindromic CRE site from the promoter region of human somatostatin gene (5’-CC<u>TGACGTCA</u>TCCG-3’)(Unlabelled CRE, Fig. S2b). The binding of CREB to the non-target DNA was assessed with double-stranded linear 14 bp competitor oligonucleotide (5’-GGCTAAAGCATTCT-3’) with and without 5’ AlexaFluor® 488 labelling. Binding of monomeric bZIP peptides lacking the leucine zippers (bZIP-ΔLZ) were assessed with linear 14 bp double-stranded DNA labelled at the 5-terminus withAlexaFluor® 488: DNA sequence for ATF2 and MafG was 5’-CCTGACGTCATCCG-3’for Jun, JunD and Fos was 5’-GGATGAGTCATTGG-3’, and for CEBPa, CEBPb and NFIL3 was 5’-CCGGCCAATCTATT-3’. Linear oligonucleotides were annealed by mixing the sense and antisense strands at 100 µM in water, heating to 95 ºC and slowly cooling down to 4 ºC over approximately an hour. Concentrations of oligonucleotides labelled with Alexa Fluor 488® at the 5’-terminus were determined by measuring the absorbance at 495 nm. The concentrations of unlabelled CRE and competitor DNA were estimated based on the absorbance at 260 nm, using extinction coefficients calculated from the nucleotide sequences.

### DNA binding kinetics

Kinetics experiments were conducted at 25 ± 0.1 °C using an SX20 stopped-flow spectrometer (Applied Photophysics). Binding of protein to Alexa Fluor 488®-labelled oligonucleotides was followed by monitoring the change in fluorescence intensity (or anisotropy) of the extrinsic fluorophore using an excitation wavelength of 495 nm and 515 nm long-pass filter(s). A 3 ms cutoff was applied for mixing time. Association measurements were performed under pseudo-first order conditions by rapidly mixing the DNA with an excess of protein at a 1:1 volume ratio. The DNA concentrations were held constant at either 5 nM (L318V) or 10 nM (CREB-bZIP-FD). Each kinetic trace was an average of 60-80 individual traces and was well fit by a single exponential decay function. To improve the signal-to-noise ratio of the CREB-bZIP-FD data, where binding was very rapid, the amplitude values were shared between the datasets collected at different concentrations.

This is reasonable since due to the extremely high binding affinity the amplitudes are not expected to vary with protein concentration under these conditions. Observed rate constants (*k*_obs_) for CREB-bZIP-FD were linearly dependent upon protein concentration, and *k*_+,*D*−*CRE*_was identified as the gradient of the line of best fit. Observed rate constants for CREB-bZIP-L318V did not follow a linear dependence. For the monomer-only binding pathway if a fast pre-equilibrium is assumed between monomer and CRE, and given the excess of protein over CRE, then data are expected to display a quadratic dependence on protein concentration, rather than a linear one, according to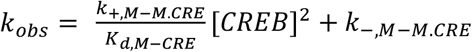. Data for CREB-bZIP-L318V were fit to a quadratic equation, constraining for physical (positive) values for all coefficients, to test this model. The independently obtained estimate for *K*_*d,M*−CRE_was used to estimate *k*_+,*M*−*M*.*CRE*_from the fitted value of the quadratic co-efficient. For the dimer-only binding pathway quantitative predictions for observed rate constants were made at many protein concentrations by numerical integration of the series of rate equations that describe the scheme in Fig. 2a (Eq. 1a-1h) in Mathematica 14 (Wolfram) (with *k*_+,M-CRE_=*k*_+,M-Comp_=*k*_+,M-M.CRE_=*k*_+,M-M.Comp_=*k*_−,M-CRE_=*k*_−,M-Comp_=*k*_−,M-M.CRE_=*k*_−,M-M.Comp_ = 0):

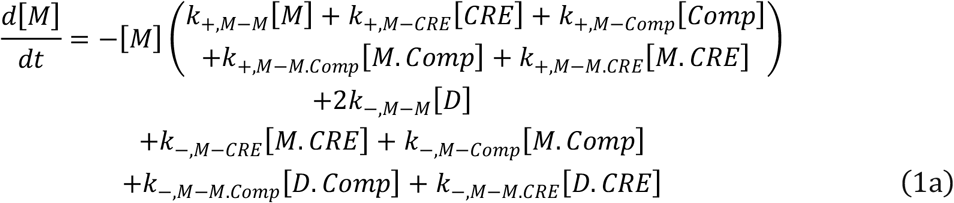

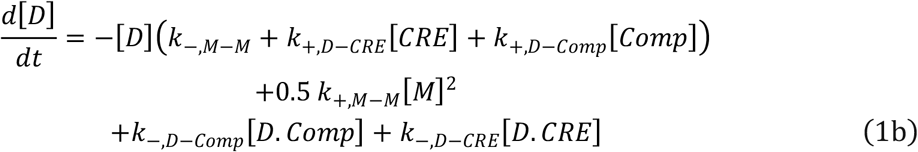

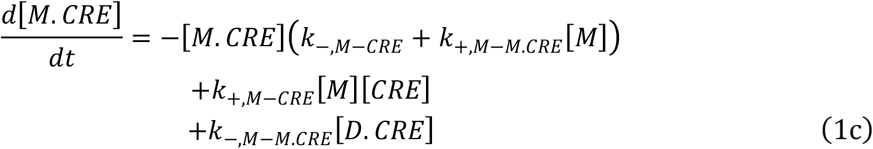

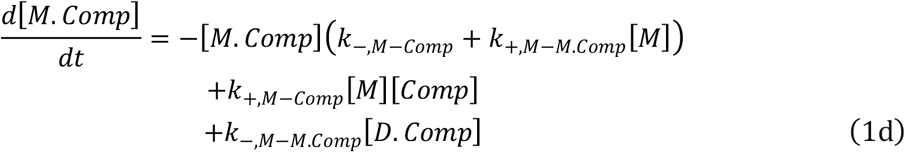

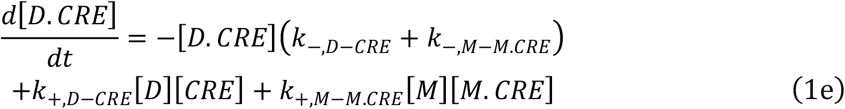

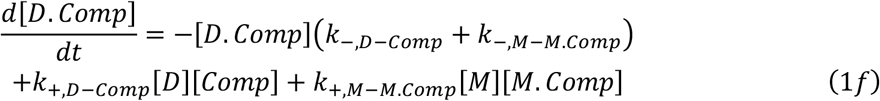

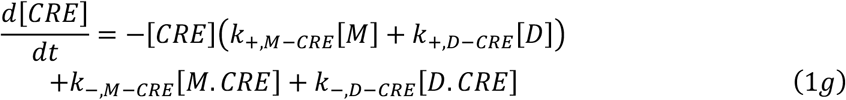

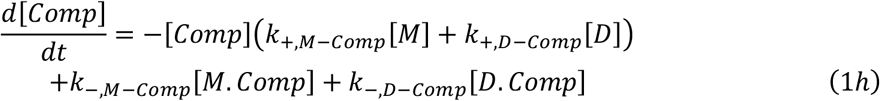

A large series of mock kinetic traces were generated for DNA concentrations matching those in the experiments, using *k*_+,*M*−*M*_ and *k*_−,*M*−*M*_ values obtained from CREB-bZIP-L318V homodimer experiments, and *k*_+,*D*−*CRE*_, *k*_−,*D*−*CRE*_, *k*_+,*D*−*Comp*_ and *k*_−,*D*−*Comp*_ values obtained from CREB-bZIP-FD DNA binding experiments. Graphs of the concentration of specific complex (D.CRE) with time were fit to single exponential decay functions to match the procedure for experimentally collected data and extract predicted rate constants.

Dissociation rate constants for CREB from AlexaFluor®488-DNA were determined in out-competition experiments by mixing the pre-formed protein-DNA complex with an excess of unlabelled CRE DNA. The concentrations of AlexaFluor®488-DNA and the protein were set at 5 nM and 100 nM, respectively. A minimum of 40 individual traces were collected for each condition, and the averages were fit to a single-exponential decay function. For CREB-bZIP-FD, the observed rate constants (*k*_obs_) were independent of unlabelled CRE concentration in the range of 1-5 µM, allowing identification of *k*_−,*D*−*CRE*_as an average of the *k*_obs_ values. The observed dissociation rate constants of the wild-type CREB-bZIP and the leucine zipper mutants were measured in the presence of 1 µM unlabelled CRE.

Dissociation rate constants for Alexa Fluor® 488-Comp (*k*_−,*D*−*Comp*_) were determined by rapid mixing of a pre-formed complex of Alexa Fluor® 488-Comp (200 nM) and CREB-bZIP (200 nM) with an excess of unlabelled CRE DNA. Since there was no fluorescence intensity change the reaction was followed by fluorescence anisotropy. A minimum of 80 individual traces were collected for each competitor concentration. The observed rate constants (*k*_obs_) were independent of unlabelled CRE concentration in the range of 0.5-2.5 µM, and *k*_−,*D*−*Comp*_ was calculated as an average of the *k*_obs_ in this range. The value of *k*_+,*D*−*Comp*_ was then calculated as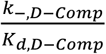.

### Equilibrium binding titrations

A series of samples containing Alexa Fluor® 488-DNA and varying amounts of protein were prepared. The concentration of Alexa Fluor® 488-DNA was at 1 or 5 nM depending upon binding affinity. After a 60 s equilibration, the samples were assayed using either a FluoroMax-4 (Horiba) or LS-55 (Perkin Elmer) spectrofluorometer. The excitation and emission wavelengths were set at 495 nm and 519 nm, respectively. The binding was followed by monitoring fluorescence anisotropy, and the overall fluorescence intensities were inferred from the measurements taken under the magic angle polarization conditions. To account for a difference in fluorescence intensity between the free and the bound states, the anisotropy readings were adjusted as described by using Eq. 2 where necessary:

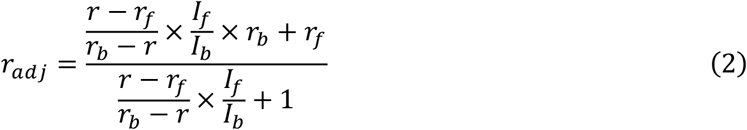

where *r*_adj_ is the adjusted anisotropy, *r* is the measured anisotropy, *I* is the measured intensity, and the subscripts *f* and *b* denote that the value is measured for the free and the bound protein, respectively.

With only two exceptions equilibrium binding titrations were analysed according to a standard two-state model equation as in Eq. 3^79^:

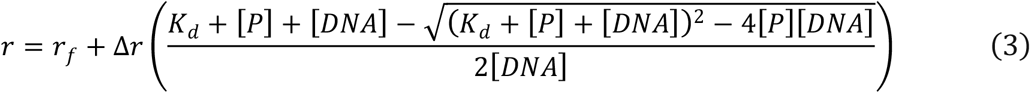

where Δ*r* is an increase in anisotropy upon binding, [*P*] is the concentration of protein, [*DNA*] is the concentration of DNA and *K*_d_ is the equilibrium dissociation constant for their binding.

Binding of dimeric CREB to Alexa Fluor® 488-Comp yielded a second transition observed at high protein:DNA ratios, necessitating fit to a three-state model equation:

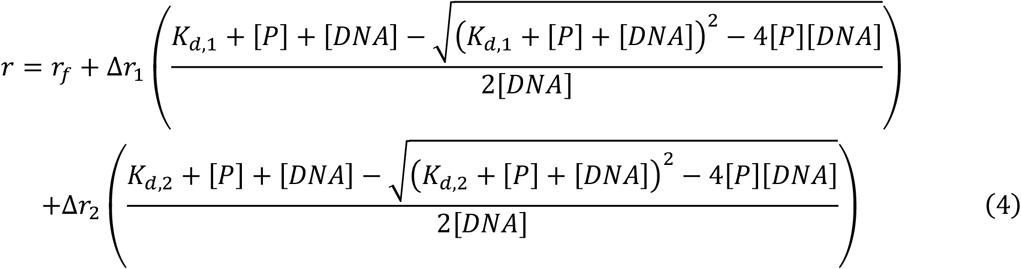

where the subscripts *1* and *2* indicate that the value characterises the first or second transition, respectively.

The DNA binding curves for the leucine zipper mutants were analysed according to a model assuming that CREB binds to DNA as a pre-formed dimer:

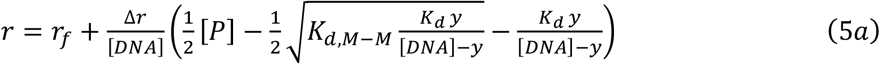

Where

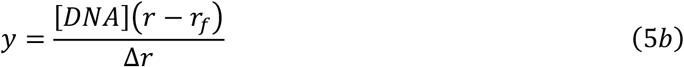

and *K*_*d,M*−*M*_ is the dissociation constant for protein dimerization. Eq. 5 is described in detail in the Supplementary Materials.

The binding of proteins to the competitor DNA was assessed by means of equilibrium competition experiments. A series of samples containing 100 nM Alexa Fluor® 488-Comp, 500 nM protein and 0-10 µM of unlabelled competitor DNA were prepared. The data was fit to Eq. 6 to extract the *IC*_50_ value:

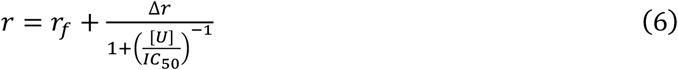

where [*U*] is the concentration of unlabelled competitor. The *IC*_50_ allows to calculate the equilibrium binding constant for the unlabelled competitor as described in^79^.

All data fitting was performed using nonlinear least squares regression analysis implemented in Prism 8-10 (GraphPad).

### Calculation of species concentrations at equilibrium

There are eight species with concentrations to be determined, requiring eight independent simultaneous equations for a unique solution. There are seven equilibrium equations for the kinetic scheme depicted in Fig. 2a, however only five are independent due to the presence of two thermodynamic squares. For convenience the five equilibrium constants that are experimentally determined are chosen (Eq. 7a-7e):

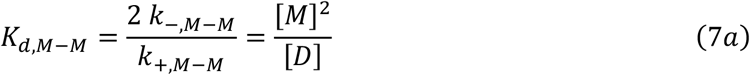

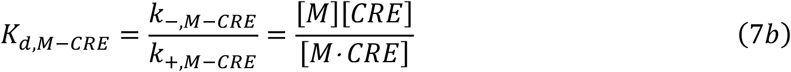

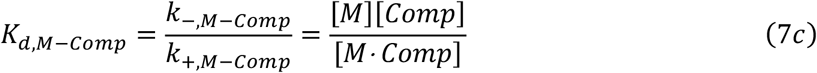

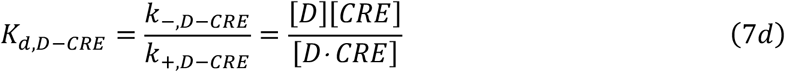

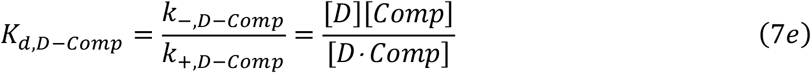

The remaining three simultaneous equations (Eq. 8a-8c) come from maintaining the total concentrations of CREB protein (*P*_T_), target CRE DNA (*CRE*_T_) and competitor DNA (*Comp*_T_):

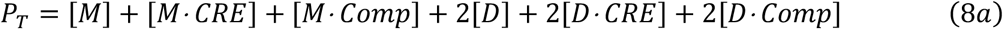

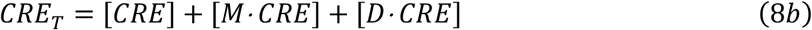

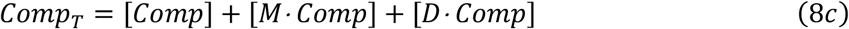

Eq. 7a-7e and 8a-8c were solved simulatenously for given total concentrations of protein, target DNA and competitor DNA using Mathematica 14 (Wolfram).

### Derivation of flux through the monomer pathway at equilibrium

At equilibrium, the proportion of target complex (D·CRE) formed through the monomer pathway (φ_M_) is:

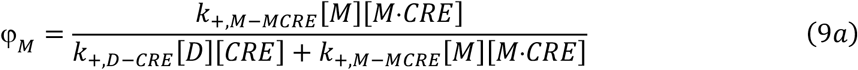

Which may be rewritten as:

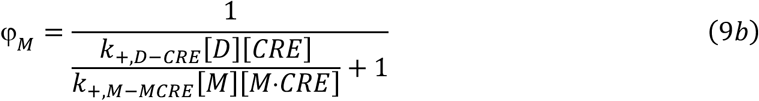

Eliminating the concentrations using Eq. 7a and 7b and simplifying gives:

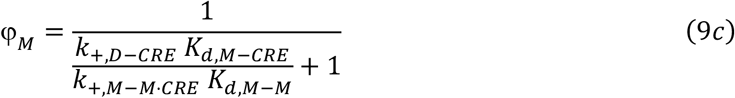

Assuming that both association reactions are very rapid and diffusion-limited i.e. *k*_+,*D*−*CRE*_~*k*_+,*M*−*M*·*CRE*_ leads to:

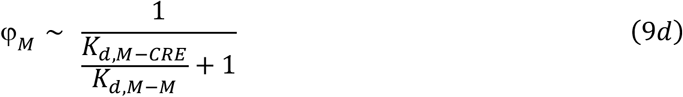

In the limiting case where *K*_*d,M*−*CRE*_≫ *K*_*d,M*−*M*_ (as we show for CREB) this further simplifies to

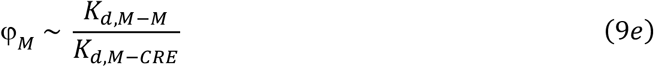

### Kinetic modelling of species concentrations and pathway fluxes

A numerical solution to the series of ordinary differential equations was found (up to 10,000 s) using Mathematica 14 (Wolfram) for a range of initial concentrations and rate constant values described in Fig. 4. Concentrations for all species at a range of timepoints from 0.001-10,000 s (organised on a log scale) were extracted from the interpolating function. The total concentrations of protein, DNA and competitor remained constant within fourteen significant figures for all solutions. At 10,000 s the concentrations of all species were as expected according to equilibrium modelling. Fluxes for complex formation by the monomer pathway and dimer pathway were calculated as *k*_+,*M*−*MCRE*_[*M*][*M*·*CRE*] and *k*_+,*D*−*CRE*_[*D*][*CRE*] respectively.

### Equilibrium dissociation constants of the dimers of leucine zipper mutants

Equilibrium constants for the monomer-dimer transitions of the leucine zipper mutants were obtained by dilution titrations. To monitor the oligomeric state of CREB bZIP, we took advantage of intrinsic tyrosine fluorescence (Y307 and Y336), that was previously observed to be quenched upon the dimer formation^69^. The proteins were serially diluted into the buffer, and the fluorescence intensity was recorded. The excitation and emission wavelengths were set at the maxima of the excitation and emission spectra (278 nm and 307 nm, respectively). The concentration of L318V ranged from 0.5 to 150 µM, and the concentration of L332V ranged from 0.3 to 60 µM. The observed fluorescence intensities were normalised by the concentration of protein in the sample, and the monomer-dimer *K*_D_ was extracted by fitting the data to Eq. 10:

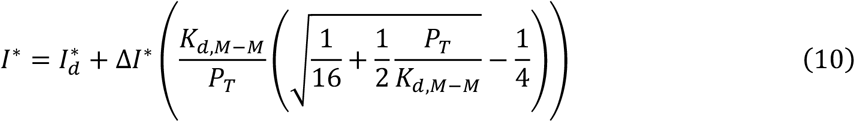

where *I*\* is the normalised fluorescence intensity of the sample, *I*_d_* is the normalised fluorescence intensity of fully dimeric protein, Δ*I*\* is a decrease in normalised fluorescence intensity upon protein dimerization and *P*_T_ is the total protein concentration. Derivation of Eq. 10 is described in detail in Supplementary Materials.

### Dissociation kinetics of the dimers of leucine zipper mutants

Concentrated protein samples (24.75 µM of L318V and 11 µM of L332V) were rapidly diluted with buffer at 1:10 ratio, and the change in intrinsic tyrosine fluorescence upon the dimer dissociation was recorded. The excitation wavelength was set to 278 nm, and the emitted light was detected after passing through a 305 nm cut-off filter. A minimum of 60 traces per mutant were collected and averaged for the analysis. The averaged traces were fit to Eq. 11a:

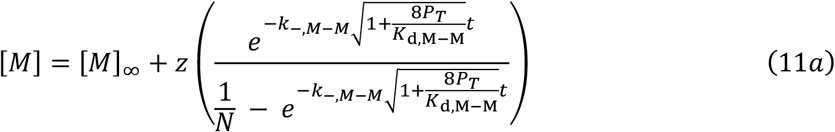

Where

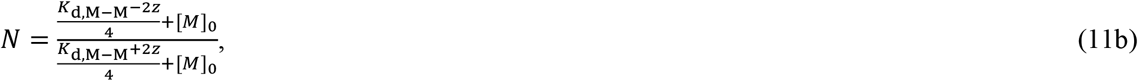

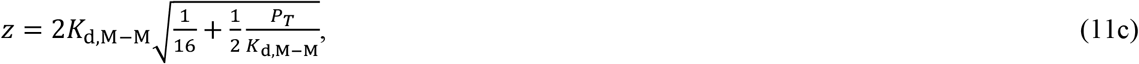

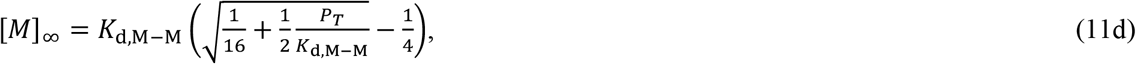

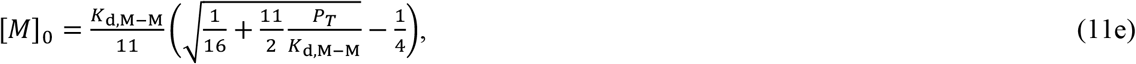

*K*_d,M−M_ is the equilibrium dissociation constant for the monomer-dimer transition, *k*_−,*M*−*M*_ is the dissociation rate constant of the protein dimer and *P*_T_ is the total protein concentration in the cell. Derivation of Eq. 11 is described in Supplementary Materials. The fitting was carried out using pro Fit (QuantumSoft) software; the script is available upon request.

### Homodimerization kinetics of wild-type CREB bZIP

Homodimer formation was monitored by following the change in intrinsic tyrosine fluorescence on an SX20 stopped-flow spectrophotometer (Applied Photophysics, Leatherhead, UK). CREB in 8 M urea was mixed rapidly in a 1:10 volume ratio with buffer solutions of various (lower) urea concentrations, to achieve protein and urea concentrations outlined in the text. An excitation wavelength of 278 nm was used with a 305 nm cut-off filter, and the temperature was maintained at 25.0°C. At least 50 traces were averaged for a typical measurement.

Kinetic refolding traces were fit using proFit (QuantumSoft, Tomsk, Russia) software to Eq. 12a

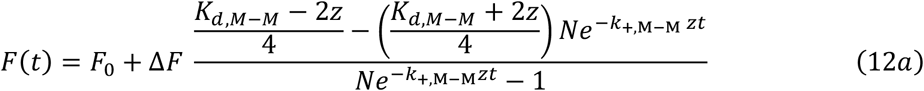

Where

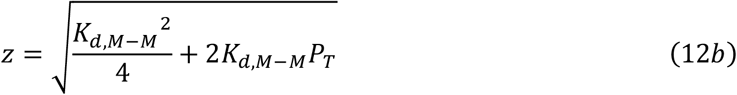

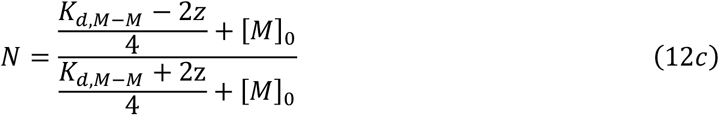

and *P*_*T*_ is the total CREB-bZIP concentration.

Due to the presence of 8M urea in the protein solution prior to mixing we assume *M*_0_ = *P*_T_. The equation is derived in^69^. *K*_*d,M*−*M*_ without urea was extracted as the y-intercept of a straight line fit of *K*_*d,M*−*M*_ against urea concentration in Fig. 2b. The value of *k*_+,M-M_ in the absence of urea was extracted as the y-intercept of a straight line fit of *k*_+,M-M_ against urea concentration in Fig. 2c.

## RESULTS

### Development of a kinetic scheme for target search

Within the cell, transcription factors must locate their cognate sites amongst a large excess of alternative sequences. We aimed to reveal the stoichiometry of the target search process of a disordered DNA binding domain, using a simplified in vitro experimental system that allowed for maximal control of variables such as concentration. We chose to examine the bZIP domain of CREB: CREB operates largely as a homodimer^38^, and its binding sites are identified and quantified^80^. It is expressed at similar levels in all cell types (Human Protein Atlas proteinatlas.org) ^81,82^ and the number of molecules per cell has been quantified in two studies with different techniques and cell types^83,84^.

These features make it well suited for downstream modelling purposes, simplifying analysis and extrapolation to cellular context. The dominant pathway is determined by kinetic considerations and may not simply reflect the concentrations of free monomer and dimer within the cell. Within the nucleus, the cognate DNA binding sites are also concealed within a vast excess of non-target DNA which will alter the balance between the species, and it is not immediately obvious how this will impact pathway preference. In principle both monomer and dimer pathways take place, with the values of the rate constants and the protein and DNA concentrations controlling the rate of complex formation (flux) through each pathway. Thus, we considered non-target DNA in our experiments and modelling to examine its importance. To limit the complexity of the system the DNA target was placed in a short double-stranded oligonucleotide hairpin. The simplest model suitable for describing such a system is outlined in Fig. 2a. In this model CREB molecules can dimerise, and can bind to both target DNA and non-target (competitor) DNA in reversible reactions. In this simplest possible consideration, there are already at least six different distinct states the protein can exist in, and at least seven different simple binding reactions. Importantly the scheme makes no assumptions about the search stoichiometry, but rather this information will naturally emerge from the consideration of the rate constants and concentrations once they are established. We aimed to experimentally assign values to as many of the equilibrium and rate constants in Fig. 2a as possible.

Formation and dissociation of CREB homodimers was probed in stopped-flow kinetic urea refolding experiments. A bZIP domain corresponding to residues 277-339 of CREB-A, hereafter described as CREB-bZIP, was first incubated in 8 M urea to unfold the protein, and then rapidly mixed with buffer of varying (lower) concentrations of urea. Upon the ensuing rapid decrease of urea concentration the population of dimeric (partially folded) CREB increased, resulting in a change in intrinsic tyrosine fluorescence. Fitting individual kinetic traces with a reversible model (centre line, Fig. 2a) allows a simultaneous extraction of estimates for both kinetic rate constants for folding (*k*_+,*M*−*M*_) and equilibrium dissociation constant for binding (*K*_d, M-M_) at each urea concentration. Extrapolation of a straight-line fit back to 0 M urea then further estimates *k*_+,*M*−*M*_ and *K*_d, M-M_ in the absence of denaturant to be (10 ± 2) μM^−1^s^−1^ and (5.7 ± 0.3) nM (Fig. 2b-c, Table S1).

We moved on to describe the reaction between dimeric CREB and its target DNA CRE. To generate a fully dimeric variant, a cross-linked version of CREB-bZIP was generated. This was done by the addition of three amino acids at the C-terminus (GGC) which oxidised to form disulphide-linked “forced” dimer (CREB-bZIP-FD) (Fig. S2a). Varying concentrations of CREB-bZIP-FD were incubated with 1 nM Alexa Fluor® 488-labelled self-annealing oligonucleotide (Fig. S2b) containing the CRE sequence (termed Alexa Fluor® 488-CRE). Upon protein binding the probe fluorescence intensity decreased, and anisotropy increased. Equilibrium curves indicated a sub to low-nM upper estimate for the equilibrium dissociation constant with Alexa Fluor® 488-CRE (*K*_d, D-CRE_) (Fig. S3).

The association kinetics with Alexa Fluor® 488-CRE were examined under pseudo-first order conditions using fluorescence stopped-flow spectroscopy. Kinetic traces fit well to single-exponential decay functions (Fig. S4), characterised by an apparent rate constant that was linearly dependent upon the bZIP concentration (Fig. 2d). Therefore, it was possible to identify the gradient of the straight-line fit (2.57 ± 0.06 nM^−1^s^−1^) as the association rate constant for dimeric CREB bZIP with CRE (*k*_+,D-CRE_). The CREB:CRE interaction is thus extremely rapid; such a high value is indicative of significant electrostatic rate enhancement. Kinetics of dissociation were next examined using a traditional out-competition assay, by rapid mixing of pre-formed complexes of Alexa Fluor® 488-CRE DNA and CREB CREB-bZIP-FD with large 20 to 100-fold excesses of unlabelled CRE DNA. Kinetic traces were again well fit by single-exponential decay functions (Fig. S5a), and the apparent rate constants did not depend upon the concentration of unlabelled CRE (Fig. 2e). The average apparent rate constant, identified as the dissociation rate constant for dimeric CREB bZIP from CRE (*k*_−,D-CRE_) was 0.556 ± 0.008 s^−1^. *K*_D, D-CRE_ was then estimated as *K*_D, D-CRE_ = *k*_−,D-CRE_/*k*_+,D-CRE_ = 214 ± 6 pM for CREB-bZIP-FD, consistent with the limit provided by equilibrium titration. In support of this approach, the directly determined (2.01 ± 0.15 nM) and kinetic estimate (1.84 ± 0.05 nM) for *K*_D, CRE_ of a weaker binding mutant (R284_K285insG_3_) are the same within error (Fig. S3). Overall, the rate and equilibrium constants characterising the interaction for CREB-bZIP-FD with CRE demonstrate the high kinetic and thermodynamic stability of the complex.

We next sought to characterise the interaction of CREB-bZIP-FD with non-target DNA sequences. We utilised short dsDNA oligonucleotides to limit multiple CREB dimers binding on the DNA and facilitate accurate calculation of equilibrium dissociation constants for modelling purposes.

Equilibrium titration experiments were repeated using a fluorescently-labelled 14 base pair double-stranded competitor oligonucleotide (Alexa Fluor® 488-Comp, Fig. S2b), designed to lack CRE half- and full-sites. Two binding transitions associated with an increase in fluorescence anisotropy were observed (Fig. S6a); the second transition occurs at very high protein:DNA ratios not expected within the cell and may represent either the binding of two CREB dimers to the DNA or the formation of a DNA-bound CREB tetramer. Dimeric CREB binds fairly tightly to Alexa Fluor® 488-Comp DNA, with the lower *K*_D,D-Comp_ being in the hundreds of nM (Fig. 2f, Table S2). This still indicates an approximately three orders of magnitude specificity for CRE DNA recognition. By suitable arrangement of conditions, it was possible to observe the dissociation process using fluorescence anisotropy stopped-flow (Fig. S5b). Using this approach, we were able to estimate the dissociation rate constant (*k*_−,D-Comp_) for CREB-bZIP-FD from Alexa Fluor® 488-Comp as 158 ± 3 s^−1^ (Fig. 2g, Table S2). Assuming a two-state reaction, we infer an association rate constant *k*_+,D-Comp_ = *k*_−,D-Comp_/*K*_D, D-Comp_ = 1.0 ± 0.1 nM^−1^s^−1^, which is very similar to that obtained with target DNA in association experiments. Thus, binding rate constants for dimeric CREB appear relatively insensitive to DNA sequence.

The remaining uncharacterised reactions in Fig. 2a all involve CREB monomer binding to DNA, so a forced monomeric variant was designed. Since the variant would exist in the presence of very high DNA concentrations in some assays we reasoned that as much of the LZ as possible should be removed to negate any templated homodimerization. The helical prediction software AGADIR indicated that CREB 277-315 (CREB-bZIP-ΔLZ) should contain the same residual helical content within the basic region as a full-length CREB monomer (Fig. S1). Unfortunately full kinetic characterisation of the scheme in Fig. 2a was hindered since it was not possible to monitor the kinetics of reactions between CREB-bZIP-ΔLZ and DNA. Equilibrium titration indicated the monomeric CREB CREB-bZIP-ΔLZ bound only very weakly to Alexa Fluor® 488-CRE, with an affinity in the hundreds of μM (Fig. S6b, Fig 2f, Table S2) i.e. the binding affinity is reduced by almost six orders of magnitude compared to the dimeric CREB-bZIP, despite no DNA-contacting residues having been removed. In equivalent equilibrium titration experiments CREB-bZIP-ΔLZ bound Alexa Fluor® 488-Comp very weakly as well. Indeed the binding affinity is remarkably similar to that for target DNA (Fig. 2f, Table S2) indicating an extremely poor target specificity from a thermodynamic perspective. This weak target binding of the monomer not only impedes its kinetic characterisation, but as we shall demonstrate has significant implications for the efficacy of a monomer binding pathway for target binding.

### Modelling highlights dimeric CREB as the active species at equilibrium

Considering nuclear CREB concentrations of around 1 µM^83^ and a *K*_D, M-M_ in the tens of nanomolar, one can naively suggest CREB is mostly dimeric in the nucleus. However vast excesses of non-target DNA may provide a template for dimerization, or compete for monomer binding. The model (Fig. 2a.) allows us to extrapolate and make predictions for such high DNA concentrations since five of the seven equilibrium constants have been determined. The remaining two equilibrium constants can be indirectly calculated by identifying the presence of two thermodynamic cycles within the reaction scheme. Thus, we can use the model to estimate equilibrium concentrations of all species for any given mixture of protein, target and competitor DNA by solving its associated series of simultaneous equations (Eq. 7a-7e and 8a-8c)(Fig. 3). To provide a model of the Eukaryotic nucleus we estimated nuclear concentrations by considering reports of the number of CREB molecules and CREB target sites^80,83^ and typical nuclear dimensions (1 µM and 0.1 µM, respectively). For perspective, concentrations of around 1 mM of the competitor provide base pair concentrations roughly equivalent to those present in the nucleus. We performed modelling at this concentration and a range of significantly lower concentrations to account for significant uncertainty in how much of this DNA might be accessible in the nucleus. The modelling demonstrates clearly that the total dimeric fraction of CREB is expected to increase with competitor concentration. For competitor concentrations above 1 µM the vast majority of CREB is expected to be found in competitor-bound CREB dimers. This DNA concentration is roughly three orders of magnitude lower than expected within the Eukaryotic nucleus. The modelling reveals the least populated species is CRE-bound monomer, which has concentrations many orders of magnitude below all other protein states (including CRE-bound dimer).

**Fig. 3.**
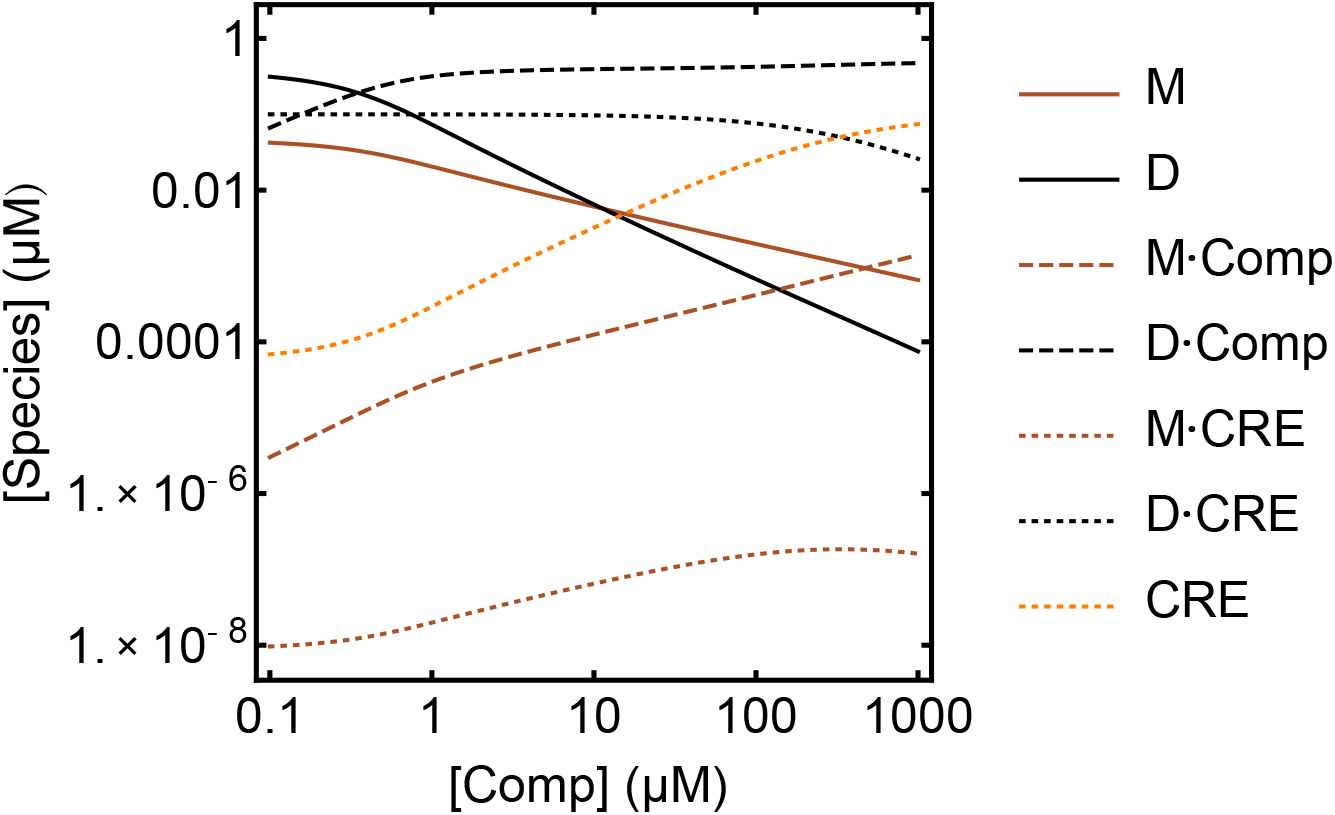
Modelling of equilibrium concentrations of species in mixtures of protein, target CRE DNA and competing non-target DNA. Species shown are free monomer (brown solid line), free dimer (black solid line), free target (orange dotted line), competitor bound monomer (brown dashed line), competitor-bound dimer (black dashed line), CRE-bound monomer (brown dotted line) and CRE-bound dimer (black dotted line). Concentrations of CREB and CRE are 1 µM and 0.1 µM respectively, chosen to approximate those found within the eukaryotic cell nucleus. Equilibrium constants used for modelling are those calculated in Fig. 2 (Table S2) for CREB-bZIP (*K*_d,M-M_ = 5.7 nM, *K*_d,M-CRE_ = 210 µM, *K*_d,M-Comp_ = 440 µM, *K*_d,D-CRE_ = 214 pM, and *K*_d,D-Comp_ = 158 nM). Modelling is shown for 14 base pair (bp) competitor DNA concentrations ([Comp]), from 0.1 to 1000 µM, equivalent to base pair concentrations of 1.4 to 14,000 µM. Above 10 µM competitor DNA most free CREB is monomeric, however most CREB is dimeric, with the majority species being competitor-bound dimer. Importantly monomer-bound CRE is barely populated under all conditions.

**Fig. 4.**
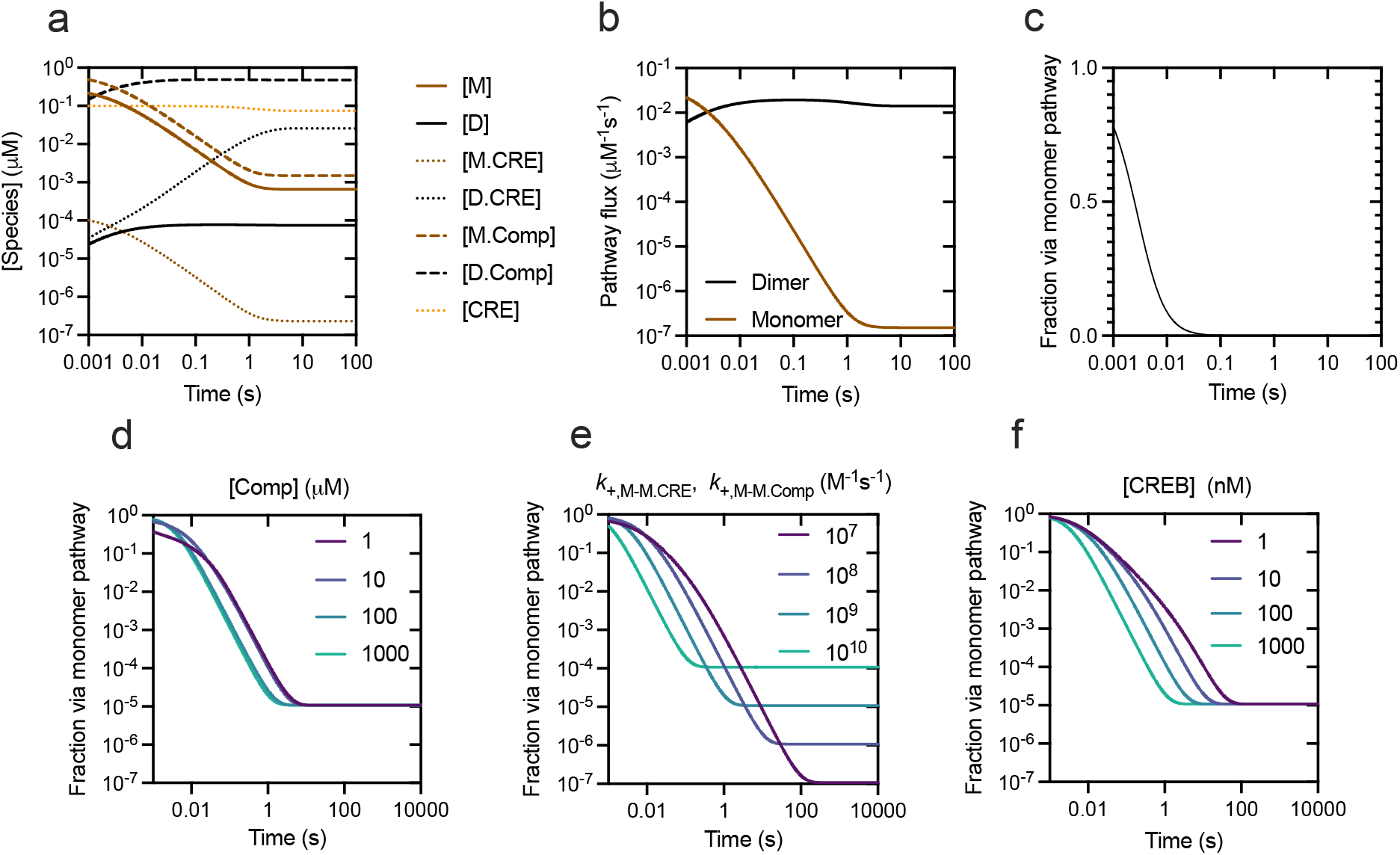
Kinetic modelling of species concentrations in mixtures of CREB and DNA demonstrates preference for dimer pathway is established quickly. Initial conditions are chosen to favour the monomer pathway with all CREB being monomeric. Unless otherwise stated in the legends concentrations and rate constants are as follows: [CREB] = 1 µM, [CRE] = 0.1 µM, [Comp] =1000 µM (chosen to approximate those found within the eukaryotic cell nucleus), *k*_+,M-M_ = 10 µM^−1^s^−1^, *k*_−,M-M_ = 0.029 s^−1^, *k*_+,D-CRE_ = 2570 µM^−1^s^−1^, *k*_−,D-CRE_ = 0.55 s^−1^, *k*_+,D-Comp_ = 1000 µM^−1^s^−1^, *k*_−,D-Comp_ = 158 s^−1^, *k*_+,M-CRE_ = *k*_+,M-Comp_ = *k*_+,M-M.CRE_ = *k*_+,M-M.Comp_ = 1000 µM^−1^s^−1^, *k*_−,M-CRE_ = 210 µM *k*_+,M-CRE_, *k*_−,M-Comp_ = 440 µM *k*_+,M-Comp_. Values for *k*_−,M-M.CRE_ and *k*_−,M-M.Comp_ were set as required to maintain thermodynamic cycles. Evolution of species concentration (**a**), pathway flux **(b**) and fractional flux through the monomer pathway (**c**) for these values of variables demonstrate that the dimer pathway quickly overtakes the monomer pathway in terms of flux, and in advance of appreciable population of the specific D.CRE complex. Rapid collapse to the dimer pathway is observed over a large range of competitor concentrations (**d**), association rate constants for monomer with monomer-bound DNA (**e**) and CREB concentrations (**f**).

We then considered the fluxes throughout the scheme that maintain these equilibrium conditions. The proportion of CRE binding through the monomer pathway is independent of total concentrations of CREB, CRE and competitor, and can be approximated under our conditions as 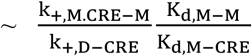 (see Eq. 9a-9d, Materials and Methods). We have no experimental estimate of *k*_+,M·CRE-M_, however it likely lies somewhere between the rate constant for homodimerization (*k*_+,M-M_ ~ 10^7^ M^−1^s^−1^) and that for dimer-CRE binding (*k*_+,D-CRE_ ~ 10^9^ – 10^10^ M^−1^s^−1^). Rate constants in the range 10^9^ – 10^10^ M^−1^s^−1^ are typically described as diffusion-limited reactions and very rarely exceeded. Given the electrostatically favourable nature of the interaction, and the locally high concentration of the two bZIP monomers in the encounter complex with the small DNA construct, values in this upper range appear most likely. Even if this extremely high upper limit is considered, this would indicate that only one in roughly 10^4^ CRE complexes is formed through the monomer pathway i.e. at equilibrium the dimer search pathway is strongly favoured over the monomer pathway for CREB.

### Equilibrium conditions are approached rapidly

The relative flux through the two pathways will be different before equilibrium is established, so we investigated the anticipated timescales for approach to equilibrium and expected relative flux during that period. Only six of the rate constants could be determined experimentally (black in Fig. 2a) precluding full unambiguous kinetic modelling of the entire scheme. However, if the remaining association rate constants are estimated, then all rate constants may be inferred and species concentrations modelled with time for set starting conditions using numerical integration of the rate equations. Previous work with the GCN4 bZIP demonstrated that monomeric and dimeric GCN4 bound target DNA with similar association rate constants^41^. Therefore, we estimated *k*_+,M-CRE_ and *k*_+,M-Comp_ as 10^9^ M^−1^s^−1^ to match the approximate values we obtained for *k*_+,D-CRE_ and *k*_+,D-Comp_, which were very similar to each other and in the range typically described as diffusion-limited. Values for *k*_−,D-CRE_ and *k*_−,D-Comp_ were then inferred from the relevant equilibrium dissociation constants. We performed modelling for a range of rate constants for *k*_+,M·CRE-M_ and *k*_+,M·Comp-M_ (10^7^ – 10^10^ M^−1^s^−1^). Using these physically sensible values as estimates demonstrates the system is expected to approach equilibrium rapidly, and in useful biological timescales of under one second (Fig. 4). We considered factors that would slow the approach to equilibrium. To deliberately favour the monomer pathway all CREB was assumed to start as entirely unbound and unfolded monomer. Decreasing CREB concentration extends the time taken to reach equilibrium as expected, however 99% of flux occurs through the dimer pathway in under 0.5 s even when the CREB concentration is only 1 nM (1000-fold lower than estimated in the nucleus) (Fig. 4f). Some non-target DNA in the nucleus contains sequences similar to the CRE consensus or half-site which would sequester the CREB dimer for longer. This should not impact the pathway preference at equilibrium; however it would reduce the occupancy of target sites and delay the approach to equilibrium. Our modelling suggests the delay is not significant - even if we allow the competing DNA to be a highly unrealistic maximally effective kinetic trap (by matching *k*_−,D.Comp_ to *k*_−,D.CRE_), most flux is still via the dimer pathway by around 50 ms (Fig. S7).

In all modelled scenarios the dimer pathway reaches parity with the monomer pathway in under 100 ms and then begins to dominate. Thus, systems containing physiologically realistic concentrations of CREB and DNA approach equilibrium control, where the dimer pathway is strongly favoured, very rapidly.

### Target search efficiency is controlled by non-target affinity

A key implication from the modelling is that in the presence of excess competitor DNA most dimeric protein will be bound non-specifically. If we assume a dimeric search pathway, characterised by only the three reversible reactions that we have measured kinetically we can quantitatively predict the observed rate constants for dimeric CREB with CRE in the presence of excess competitor.

Importantly, this may be tested experimentally. We repeated the kinetic association experiments in the presence of one to four µM unlabelled competitor DNA (Fig. S2b), which corresponds to a concentration of 15 - 60 µM in terms of DNA base pairs. As anticipated, the observed rate constant was successively reduced with increasing concentrations of competitor (Fig. S8, Fig. 5a). The values of *k*_+,D-Comp_ and *k*_−,D-Comp_, combined with the excess concentration of competitor DNA ensure that a fast pre-equilibrium is established between CREB dimer and competitor DNA, so the model predicts a linear relationship between observed association rate and free dimer concentration. The extent of observed “slow-down” in specific binding should be equal to the proportion of CREB dimer that is free to bind. Therefore, the measured *K*_d,D-Comp_ of CREB-bZIP-FD to unlabelled competitor DNA (Fig. S9) was used to calculate the expected free dimer concentration at each DNA concentration. The “slow-down” is broadly consistent with the predicted line y=x (Fig. 5b), with a straight line fit of gradient 1.08 ± 0.04. This demonstrates the observed target search rate constants can be rationalised fully by a simple sequestration model, and are dictated by a combination of *k*_+,D-CRE_ (the fundamental rate constant for association of dimer and its DNA target), the competitor DNA concentration and *K*_d,D-Comp_ (the affinity of dimer for competitor DNA).

**Fig. 5.**
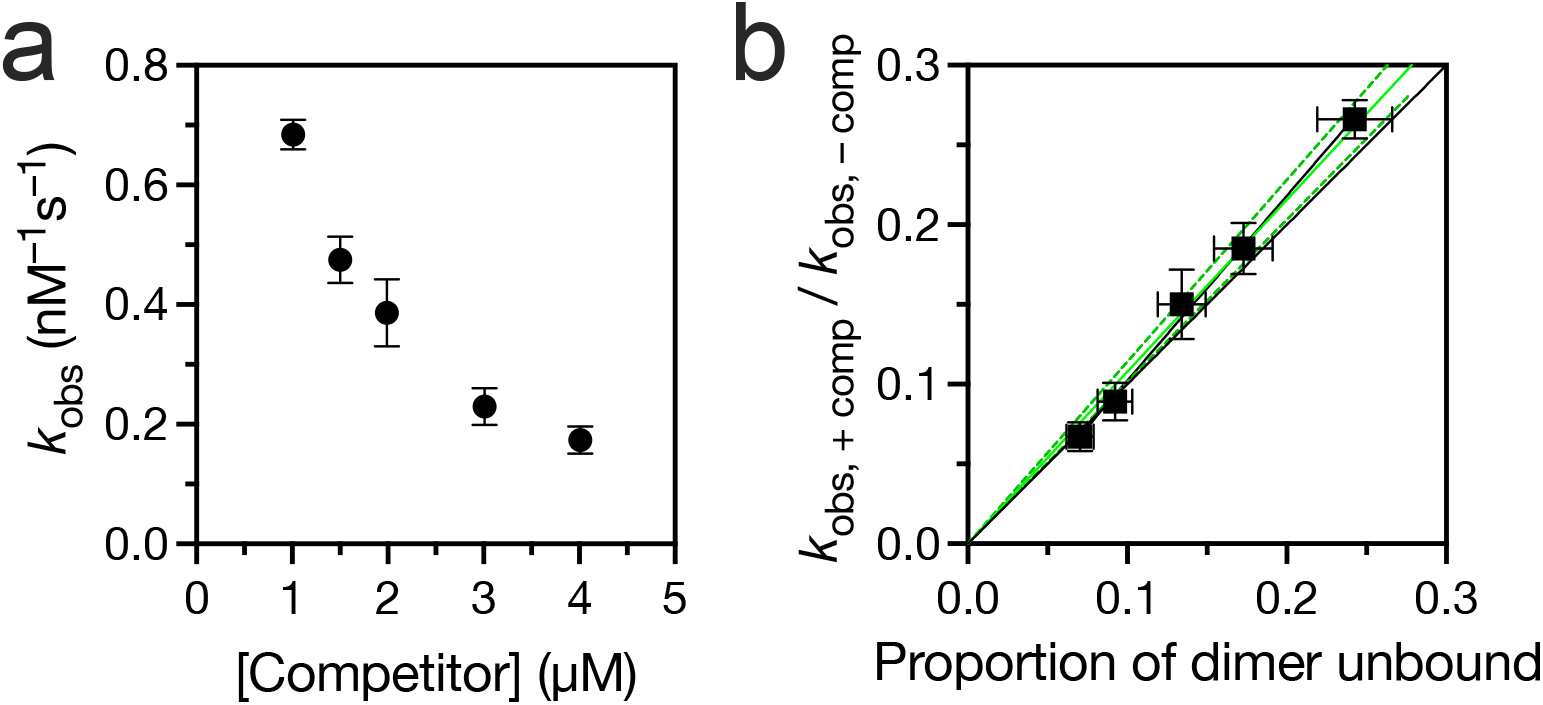
Retardation of association of CREB-bZIP-FD with Alexa Fluor® 488-CRE by competing non-target DNA is accurately predicted by the model. **a**, Observed association rate constants for 10 nM Alexa Fluor® 488-CRE DNA with CREB-bZIP-FD in the presence of varying concentrations of unlabelled competitor. Experiments were performed by diluting the protein into a mixture of Alexa Fluor® 488-CRE and unlabelled competitor DNA. Observed association rate constants were obtained from the gradients of straight-line fits for CREB-bZIP-FD binding to CRE DNA (Fig. 2d, Fig. S8). The errors are the errors of fit. **b**, Fractional decrease in observed association rate constant (relative to without competitor) plotted against the equilibrium fraction of the free protein in the reaction mixture. Errors are propagated according to standard rules for both axes. Solid black line is y = x, which represents the predicted fraction decrease if dimer sequestered by the unlabelled non-target competitor DNA is unable to bind to CRE. Solid green and dashed green lines represent the line of best fit to the data and 95% confidence intervals, respectively.

### A significantly destabilised dimer still binds DNA as a dimer

To further establish the legitimacy of our modelling approach, and therefore its conclusions, we used the model to make predictions for the results of independent kinetic experiments, starting far from equilibrium and favouring the monomeric form of CREB. We made use of reported destabilising mutations in the leucine zipper^85^, L332V and L318V (Fig. 1). The dimerization reactions of the two variants were characterised using tyrosine fluorescence as a probe; *K*_D, M-M_ was estimated using equilibrium fluorescence intensity as a function of protein concentration (Fig. 6a), and dimer dissociation rate constant (*k*_−,M-M_) estimated using a rapid dilution stopped-flow strategy (Fig. 6b).

**Fig. 6.**
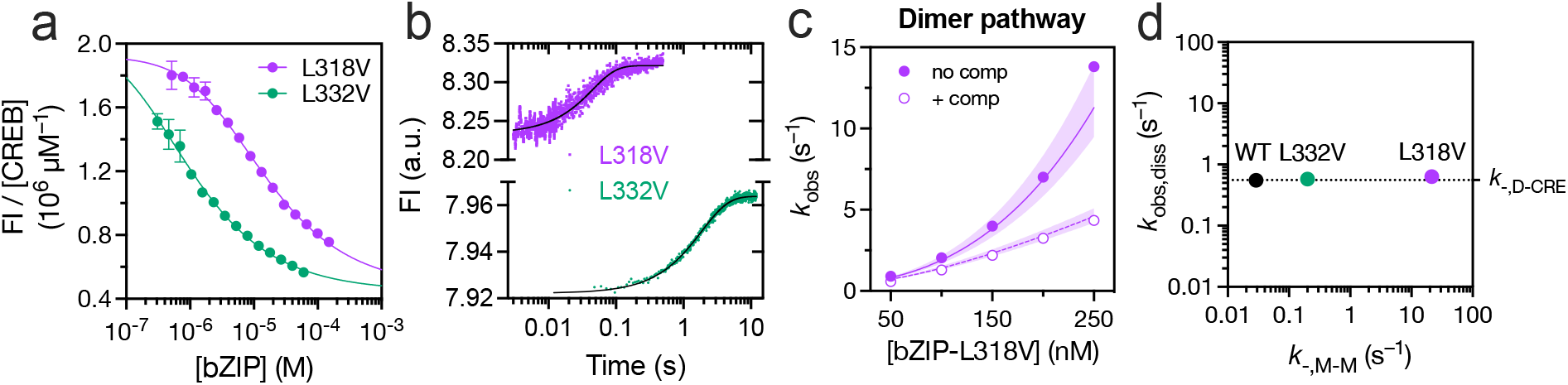
Destabilised CREB bZIP dimers still favour a dimer pathway for CRE binding. **a**, Tyrosine fluorescence dilution equilibrium curves for CREB-bZIP-L318V and CREB-bZIP-L332V. Fluorescence is corrected for the change in protein concentration between each sample. Solid lines represent lines of best fit to Eq. 10. **b**, Tyrosine fluorescence kinetic traces following 11-fold dilution to encourage dimer dissociation. Final protein concentrations were 2.25 μM for CREB-bZIP-L318V and 1.00 μM for CREB-bZIP-L332V. Black lines represent best fit to Eq. 11, where *K*_d,M-M_ is fixed at the value obtained in Fig 4a. **c**, Observed association rate constants for CREB-bZIP-L318V upon mixing with 5 nM AlexaFluor®488-CRE DNA in the presence (open circles) and absence (closed circles) of 1 μM competitor DNA. Solid and dashed lines represent predictions (c.f. fits) for a dimer-only binding model obtained using numerical integration of the rate equations, where rate constants were obtained in independent experiments (Fig 2d, 2e, 2g, 4a, and 4b). The shaded purple area represents the confidence interval for the predictions based on experimental uncertainties rate constants. **d**, Observed dissociation rate constants for the D·CRE complex (~*k*_−,D-CRE_ + *k*_−,M-M.CRE_) plotted against dissociation rate constant of the bZIP dimer (*k*_−,M-M_). The dotted line is at the value of *k*_−,D-CRE_ estimated using CREB-bZIP-FD.

**Fig. 7.**
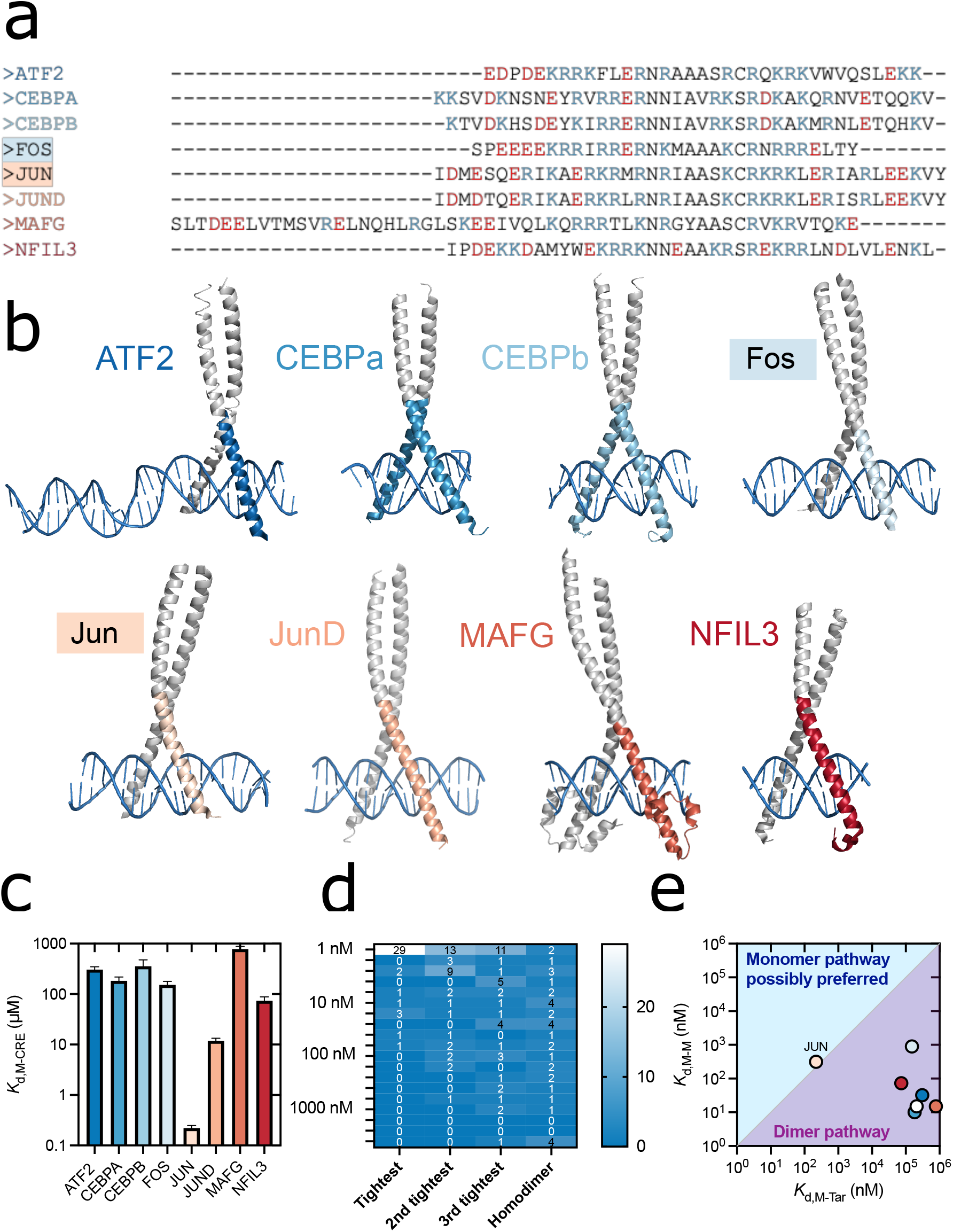
Monomeric bZIPs have generally low affinities for their DNA targets. **a**, Manually aligned amino acid sequences of peptides used. Acidic residues are coloured red, basic residues are coloured blue. Sequences were extended past the Uniprot defined N-terminal start of the basic residue to include any stabilising structures. Sequences were also extended past the Uniprot defined N- and C-terminal ends of the basic region to match predicted helicity of the full-length protein without including significant zipper interface (Fig. S13). **b**, X-ray crystal structures of ATF2 (PDB-1T2K^101^), CEBPa (PDB-8K8C^102^), CEBPb (PDB-1GU4), FOS (PDB-1FOS^103^), JUN (PDB-1FOS^103^), JUND (PDB-5VPE^104^), MAFG (PDB-7×5e^105^) and NFIL3 (PDB-8K86^102^) bZIP peptides in complex with DNA. The sequence used for each peptide is mapped in colour onto the bZIP (white). **c**, Equilibrium dissociation constants for bZIP-ΔLZ peptides with AlexaFluor®488-labelled target dsDNA oligonucleotides (*K*_d,M-Tar_) obtained from fluorescence anisotropy equilibrium binding curves (Fig. S14). Error bars indicate fitting error for the average curve from triplicate experiments. Jun has a binding affinity >50-fold higher than the next tightest binder (JunD), despite their 90% sequence identity. **d**, Heat-map analysis of equilibrium dissociation constants for dimerization (*K*_d,M-M*_) of bZIPs reported in Reinke et al. for 39 bZIP peptides from microarray experiments^38^. Data were binned using a log-scale. **e**, Comparison of *K*_d,M-Tar_ and *K*_d,M-M_ (homodimerization). CREB is displayed as a white circle.

L318V and L332V mutations both substantially destabilised the homodimer, increasing *K*_d,M-M_ roughly three and two orders of magnitude, respectively (Table S1). Equilibrium curves for binding of both leucine zipper mutants to AlexaFluor®488-CRE DNA could be fit to equations suitable for a dimer-only binding pathway to extract an estimate of *K*_d,M-M_ independently consistent with these values (Fig. S10). The substantive difference in *K*_d,M-M_ originates almost entirely from increase in *k*_−,M-M_ (Table S1). In contrast, dissociation kinetic experiments indicated that the zipper-destabilising mutations had very limited effect upon the apparent rate of dissociation of the bZIP.DNA complex (*k*_off_), both being under 115 % of the value obtained for CREB-bZIP despite *k*_−,M-M_ and *K*_D,M-M_ varying over three orders of magnitude (Fig. 6d). The wild-type had the same *k*_off_ as the forced dimer within error. The relative similarity of *k*_off_ in each case to the value obtained for the forced dimeric version (CREB-bZIP-FD) indicates that most CREB leaves CRE DNA as a dimer, even when the dimer has been significantly destabilised by mutation.

The most destabilised mutant, CREB-bZIP-L318V, provides an opportunity to make further progress with estimating rate constants. Under irreversible dissociation conditions, and under the expected condition where *k*_−,M-CRE_ >> *k*_−,M-M.CRE_, the observed dissociation rate constant of the CRE-bound dimer may be expected to approach *k*_−,D-CRE +_+ *k*_−,M-M.CRE_. Thus, for the most destabilised mutant, CREB-bZIP-L318V, a rough estimate of *k*_−,M-M.CRE_ = 0.079 ± 0.012 s^−1^ can be obtained and thereby *k*_+,M-M.CRE_ = (5 ± 2) nM^−1^s^−1^ inferred. The latter approximately matches the value for wild-type dimer association with CRE and competitor DNA and is at the upper end of the physically sensible range. From this it is possible to estimate that *k*_−,M-M.CRE_ */(k*_−,D-CRE +_+ *k*_−,M-M.CRE_) = 12 ± 2 % of complexes dissociate via the monomer route for the highly destabilised L318V mutant. If we assume a similar kinetic stabilisation of the dimer by DNA in both cases then we can infer only (0.017 ± 0.004) %, or roughly 1 in every 6000 complexes, dissociate via. the monomer pathway for wild-type protein.

Association kinetic experiments demonstrated that binding of CREB-bZIP-L318V to Alexa Fluor® 488-CRE DNA is significantly slower than the wild-type, with observed association rate constants no longer linearly dependent upon protein concentration. Expected values of the rate constants for a dimer-only binding pathway were obtained by numerical integration of rate equations describing the dimer-only binding pathway (black, Fig. 2a) with the previously (independently) obtained values of *k*_+,M-M_ and *k*_−,M-M_ for CREB-bZIP-L318V, and *k*_+,D-CRE_ and *k*_−,D-CRE_ from CREB-bZIP-FD association. Predictions are in excellent agreement with the experimental data (Fig. 6c). The approach also accurately predicted observed rate constants for CREB-bZIP-L318V association with CRE in the presence of unlabelled competitor (Fig 6c). The “slow-down” due to introduction of competitor depends upon protein concentration, again to the predicted extent (Fig. S11). At the lowest protein concentration of our destabilised dimer (L318V) we observe only a 1.6-fold decrease in rate constant upon introduction of competitor DNA. This demonstrates that “slow-down” due to competitor binding might not always be observed, even in the case of a dimer-only binding pathway. Thus, all our experimental target search observations for the destabilised L318V variant are also in line with expectations based on a dimer-only binding model, where dimer may be sequestered on competitor DNA.

The quality of the dimer-only binding pathway predictions may be compared with those for a monomer-only binding pathway. A quadratic dependence on concentration is expected when a fast pre-equilibrium between monomer and CRE exists according to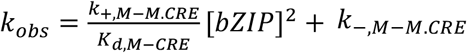. From our previous experiments we have independent estimates of all the parameters so we can predict the maximum expected rate constant for a monomer-only binding pathway (Fig. S12). These predicted values are over 10-fold lower than the observed rate constants. The data may be reasonably fit by a quadratic equation (Fig. S12), however the 95% confidence interval would imply *k*_+,M-M.CRE_ of 30 - 60 nM^−1^s^−1^. Such a high value for the association rate constant would not be completely unprecedented for a highly electrostatically favoured reaction: however the ionic strength is relatively high so significant electrostatic screening is expected, and it is over an order of magnitude higher than that already estimated so it is highly unlikely.

Taken together, the various experimental data and modelling are consistent with CREB generally arriving and leaving its DNA target as a dimer, rather than as a monomer.

### Other monomeric bZIP domains have low affinity for target DNA

The importance of the ratio of *K*_D,M-M_/*K*_D,M-CRE_ in determining the flux through monomer and dimer pathways under equilibrium conditions led us to consider *K*_D,M-CRE_ for other bZIP systems. We considered eight additional bZIPs for which structural data for the bound complex were already deposited in the PBD to inform construct design (ATF2, CEBPa, CEBPb, Fos, Jun, JunD, MafG and NFIL3). Peptide sequences (Fig. 6a) were designed to include any resolved structure (Fig. 6b) and have similar predicted helicity to the full-length protein (Fig. S13), but lack most of the leucine zipper to prevent dimerization. Seven of the monomeric bZIP-ΔLZ domains had equilibrium dissociation constants for their targets (*K*_D,M-Tar_) in the tens to hundreds of micromolar (Fig. 6c, Fig. S14). Jun bZIP-ΔLZ, is a clear outlier in the analysis: its *K*_D,M-Tar_ (220 ± 30 nM) is two to three orders of magnitude lower than the other bZIPs. However, in general, the other bZIP-ΔLZ domains behaved similarly to CREB in CRE binding experiments, displaying only very weak binding to target DNA. To interpret this finding in terms of pathway preference the *K*_D,M-Tar_ should be compared with estimated *K*_D,M-M_ (Eq. 9). For this purpose we utilised a dataset for bZIP *K*_D,M-M_ obtained previously by protein array methods by Reinke *et al*.^38^. In support of this approach the *K*_D,M-M_ reported for CREB in this dataset of 15 nM is in reasonable agreement with that obtained here from kinetic stopped-flow approaches with unlabelled CREB bZIP in solution (~ 6 nM). Except for Jun, all bZIPs examined have *K*_D,M-Tar_ several orders of magnitude higher than their *K*_D,M-M_ (Fig. 6e). Homodimer binding affinities vary significantly between bZIPs (Fig. 6d), and bZIPs may function as homodimers or heterodimers exclusively, or as both. We considered that in the case of exclusive heterodimers a more appropriate comparison for *K*_D,M-Tar_ might be with the equilibrium dissociation constant for heterodimer formation (*K*_D,M-M*_) instead of that for homodimer formation. All 38 bZIPs examined by Reinke *et al*. had at least one dimer interaction with a *K*_D,M-M*_ of under 50 nM, and values for the bZIP proteins we examined are compared with target binding affinities in Fig. S15. As these dimerization constants are all similar or lower to that for homodimerization, their comparison to *K*_D,M-Tar_ would lend even stronger support to a dimer pathway preference. In summary, following the reasoning applied for CREB, at equilibrium “active” DNA complexes for seven of eight of the bZIP DNA binding proteins would be almost exclusively generated through the dimer pathway.

## DISCUSSION

### At cellular concentrations CREB is largely dimeric

Our modelling shows for typical nuclear CREB concentrations that the dimeric state is heavily favoured over a large range of non-target DNA concentrations. Consistent with this, we interpret previous studies of CREB in cellular environments as supporting most, or all, of CREB being dimeric within the cell^86,87^. Firstly co-expression of fluorescent fusions of CREB1 activator and repressor indicates high concentrations of heterodimers by fluorescence cross correlation spectroscopy^87^. Our analysis of their results suggests 40-50% of CREB1 is present as a heterodimer since the cross amplitude falls roughly halfway between positive and negative controls. If all CREB1 was dimeric this figure would be expected to be 50% (assuming equal *K*_d,M-M_ for the two versions). Secondly, when fluorescently labelled Halo-CREB was expressed at low quantities compared with endogenous CREB in Neuro2a cells, 11% of observed CREB spots were found to dual photobleach^86^. Analysis of the Western blotting data (personal communication with the authors) indicates a maximum of (5 ± 8) % of the CREB could be fluorescently labelled. Thus, their results are consistent with 85-100% being dimeric (within one standard deviation).

### The system is characterised by similar association rate constants

The binding reaction between CREB dimer and target DNA is characterised by a very high association rate constant (~3 × 10^9^ M^−1^s^−1^). This is very similar to the value we previously obtained for wild-type CREB with a shorter variant^69^. Remarkably interactions between CREB dimer and non-target DNA, and CREB monomer and monomeric CREB.CRE complexes are also characterised by association rate constants (*k*_+_) in this same range (~1 × 10^9^ M^−1^s^−1^ and ~ 5 × 10^9^ M^−1^s^−1^ respectively).

Previously Cranz *et al*. observed GCN4 monomer and dimer to also share similar association rate constants with their target DNA^41^. Uniformly reactive spheres of similar size are expected to meet at this rate in solution. Most protein-protein and protein-DNA interactions with such high association rate constants are between oppositely charged partners, with a degree of electrostatic rate enhancement that offsets energetic barriers^88–90^. This means the similarities in *k*_+_ are perhaps slightly surprising given the differences in net charge between each of the pairs, however free CREB monomer and dimer both have positively charged basic regions that could help steer them towards the negatively charged DNA and negatively charged DNA-protein complexes. An interesting consequence of this feature is that the behaviour of the different bZIP systems is essentially controlled through the (two) rate constants for dimerization in the absence of DNA, and the various dissociation rate constants. These remaining rate constants could be viewed as determining the binding affinities, the specificity, the preferred pathway stoichiometry and the rapidity with which equilibrium in mixtures are approached. A further consequence of these high association rate constants is the rapidity with which mixtures of DNA and protein are expected to come to equilibrium. This is important because the argument in favour of the monomer search pathway is based on kinetic preference.

### Dimer pathway preferred for CREB in vitro

Predictions made according to dimer-only binding models for wild-type CREB bZIP and the leucine-zipper destabilised mutant L318V match the experimental data so we may fairly conclude that in our DNA mixtures the dimer pathway dominates; even for CREB concentrations 300-fold lower than the *K*_d,M-M_ there is still sufficient pre-existing dimer and additional dimer formation to rationalise the observed rate constants without considering monomer pathway flux. Encouragingly, our estimates of the proportion of CREB proceeding through the monomer pathway for formation and dissolution of the CRE-bound dimer are almost identical. This appears to match well with the principle of microscopic reversibility, which states a reaction follows the same trajectory in forward and reverse directions (under the same conditions)^91^. Consistent with our estimate of *k*_+,M-M.CRE_ for L318V being at the upper limit of what we used in modelling binding, the proportion of wild-type CREB-bZIP dissociating through the monomer pathway (1 in every 6000 complexes) is at the upper limit of what is projected by the modelling (1 in every 10,000 complexes). Considering the substantial differences in approach this agreement is remarkable and suggests we have correctly identified the scale of pathway fluxes. A microscopic reversibility argument has previously been used to support the monomer pathway proposal, since fusion peptides derived from the bZIP domains of GCN4 and ATF2 revealed a correlation between the rate constant for dissociation of the target complex and the kinetic stability of the dimer (*k*_−,M-M_)^46^. These studies involved swapping whole segments of sequences, potentially changing the preferred pathway, or creating unwanted allosteric effects. In contrast our analogous study made only single point mutations within the zipper region, so the approach is more targeted and the result robust and specific to CREB.

### Relevance in the cellular context

There are many obvious differences between our DNA mixtures and the nuclear environment. Our DNA is deliberately short to maximise the predictive powers of the model for the purpose of testing, whereas genomic DNA is present in chromosomes that are tens or hundreds of millions of base pairs long. This might impact residency times and eliminates the possibility for sliding long distances.

Sliding of transcription factors increases the effective “target size”, thereby increasing binding rates, and has been shown to occur inside bacterial cells^4,92^. It is likely limited within the eukaryotic nucleus for non-pioneer transcription factors however, which cannot bind to targets on nucleosome-wrapped DNA^4^. In this sense the experimental setup here is therefore not a bad representation of eukaryotic DNA, where small sections of target and non-target containing DNA are separated by nucleosomes, but nonetheless we have no DNA packaging or modifications in our mixtures. Target search may also be more complex in liquid droplets^93^, which CREB could be present in. CREB contains multiple disordered domains that could alter homodimerization and structural properties^71^ and are known to mediate interactions with other binding partners (coactivator proteins). Presence of other binding partners near CRE-containing promoters might serve to stabilise monomer-intermediate complexes (and thereby improve relative flux through the monomer pathway) if other components are present in the correct locations. On the other hand, increased local concentrations of CREB inside such assemblies are likely to substantially favour dimer formation which would further favour the dimer pathway.

Given these complexities we hope that our work will inspire carefully considered (and technically challenging) single-molecule fluorescence measurements performed inside living cells to directly visualise stoichiometry on target binding. Fortunately though, some indirect but relevant cellular data already exist to partially validate our modelling. If little CREB is indeed monomeric in the nucleus, as discussed above, this has clear and negative implications for the efficiency of a monomer search pathway. Furthermore single-molecule imaging data are also consistent with CREB largely leaving its targets as a dimer: within Neuro2a cells HaloTag-fused CREB previously yielded dissociation rate constant (0.35 ± 0.08 s^−1^) robust to the extremely dimer-disrupting L318V/L325V mutation (0.4 ± 0.1 s^−1^)^86^. This combined with observations that L318V/L325V leads to a significant decrease in promoter occupancy and transcriptional output, suggests dimer destabilisation reduces CREB target search efficiency and indicates the dimer pathway may also be preferred in cells.

In our kinetic modelling we deliberately attempted to favour the monomer pathway by initiating it with all CREB being monomeric. This is likely an extremely artificial setup. CREB does not suddenly appear at high concentrations within the nucleus. In fact CREB is almost exclusively found within the nucleus, where its concentrations stay broadly consistent independent of activation^94^. CREB may even be at least partially dimeric before translocation into the nucleus. For example the bHLH transcription factor Mad is recruited to the nucleus by dimerization with its binding partner Max, and therefore prior to reaching the DNA^68^. Despite this monomeric starting point, our numerical models quickly collapse towards equilibrium conditions where the dimer pathway is favoured. They do this within timescales many orders of magnitudes faster than cell lifetimes (hours to decades). Importantly and pertinently, previous pathway experiments with other bZIPs have been deliberately performed and discussed from the “starting point” of free monomer. We argue that this is an artificial situation and that modelling pathway flux at equilibrium may be more relevant for understanding biological systems. Within the nucleus transcription factor binding (and unbinding) takes place repeatedly and in response to signalling – so the protein is pre-equilibrated amongst the vast excess of non-target DNA and may not need to “wait” for dimerization. A dynamic quasi-equilibrium for transcriptional systems within the cell is reasonable. Equilibrium understandings have proven surprisingly effective for transcriptional regulation^95^. Protein concentration and target binding affinity are known to change transcriptional outputs^95^. In our equilibrium and kinetic modelling these factors both increase target occupancy in the expected fashion. Perhaps surprisingly, although the concentrations of the species certainly do depend on the total protein and competitor concentrations, modelling demonstrates the relative flux through each pathway does not. Thus, our conclusion that the dimer pathway is preferred, may be considered quite robust to the potential variation of these parameters between cell types and the proportion of genomic DNA that is inaccessibly packaged into chromosomes.

### Is CREB the exception or the rule?

Our results indicate that the monomeric protein has similar affinities for sequences containing, or without, the specific target for CREB. This is consistent with earlier findings for the bacterial Arc repressor, whose dimerization rates on target and non-target DNA were the same^96^. The monomer pathway requires two monomers at the target simultaneously, which is necessarily lowered from that for a single monomer binding according to the probability of one monomer being already bound.

Monomer binding affinities for target in the hundreds of micromolar, like we observed for CREB and most of the other bZIP-ΔLZs, could make this probability very small for most bZIPs under typical cellular and experimental concentrations. A state-based formalism for target site search kinetics recently concluded that dimerization in solution likely out-performs dimerization on DNA for this reason^97^. The dimer pathway is ultimately favoured for CREB because monomeric CREB binds considerably more weakly to target DNA than itself. Our analysis of a small group of bZIP proteins, indicates that CREB has typical values for both equilibrium constants, which suggests other bZIP proteins may display similar pathway preference. CREB is unusual in being described as largely an exclusive homodimer and most bZIPs have multiple binding partners^10,38^ that will have a range of expression levels. This complex signalling network is more difficult to predict in terms of behaviour. Some heterodimer complexes may have significantly higher *K*_d,M-M*_, which may push the pathway preference towards the monomer pathway. However, since all bZIP peptides examined bind at least one partner very tightly these combinations are generally assumed not to be biologically significant since they will be poorly populated. Thus, we consider it likely that most dimeric bZIP.DNA complexes result from dimer binding.

The notable exception in our analysis is Jun. The yeast homolog of Jun, GCN4 has also been shown to bind target DNA as a monomer with similar sub micromolar *K*_d,M-M_’s^41,98^. Previous pathway studies have focussed on these proteins^41–46,48^, and it is plausible that members of the bZIP family might perform search in different stoichiometries. However, these studies generally probed heterodimer formation rather than DNA target binding, aiming to answer the question of whether bZIP dimerization (and folding) happened prior or after DNA binding. Dimerization is undoubtedly accelerated by the presence of DNA, but this question is distinct to whether it locates its DNA target as a monomer or dimer in DNA mixtures. Furthermore, the ensemble mixing strategy confines observations to early times where significant changes in complex population take place, but as we have shown flux through the monomer pathway can initially be significant but drop rapidly as the reaction proceeds. All of these conditions favour observation of the monomer pathway. Future studies re-examining binding stoichiometry may be warranted.

### Avoiding kinetic traps

We find target search rates to be reduced by non-target binding. CREB dimer binds relatively tightly to non-target sequences, as reported previously^71^ and postulated to originate from a conserved cluster of basic residues in the N-terminus of CREB/ATF members^99^. A compelling argument has been that a monomer pathway could avoid significant kinetic trapping on dimeric transcription factors on non-target DNA^43,44^, which might be highly beneficial for search rates inside the nucleus. In experiments with Max proteins, which as bHLH proteins share characteristics with the bZIP family, Schepartz group demonstrated that forced dimeric versions were slowed in their target search by non-target DNA whereas wild-type versions were not appreciably slowed^43,44^. This was used to argue a kinetic advantage for the monomer pathway in the presence of non-target DNA. However, in all their experiments, as in ours, forced dimeric versions still find the targets faster than wild-type versions in DNA mixtures. For CREB we also observed reduced slow-down without forced dimerization using our destabilised zipper mutant. This appears to match the argument made for a monomer pathway avoiding kinetic traps, yet the observed slow-down actually quantitatively matches that predicted by a dimer-only binding model. We are unaware of any examples where forced (covalent) dimerization has resulted in slower target search.

It is possible that non-target DNA does not constitute an effective kinetic trap. In our experiments and models significant retardation of target search is observed but equilibrium is still approached very rapidly. For typical CREB concentrations appreciable target is occupied in useful biological timescales (under 50 ms) even for maximally high concentrations of non-target DNA. The nucleosomal nature of eukaryotic DNA, chromatin accessibility and modifications, may all reduce the impact of kinetic trapping in the nucleus. For example CREB did not occupy methylated CRE sites in ChiP assays^80^. We are unaware of any estimates of the proportion of DNA that might be accessible to CREB to adjust our models, however reduced accessibility should help the system reach equilibrium faster. Furthermore most nuclear DNA is not closely related to cognate sites, and such a feature may even be biologically required as roughly ¼ of CREB target genes do not have a full CRE^80^. It is possible that avoiding kinetic trapping may not be such a critical requirement for efficient search as previously thought.

## CONCLUDING REMARKS

Knowledge of the active search species can inform the focus and interpretation of fundamental target search studies, the development of transcriptional modulation strategies, and the design of specific DNA-binding oligomers. Research publications cite the monomer pathway as preferred for bZIPs and interpret their results in this context^50–58,64–68^. Examples include speculation of monomer pathway for other transcription factor families^51^, understanding the mechanism of action of viral accessory proteins^62^, asymmetry in binding motifs^65^, design of biologically functional photoswitchable peptides^59^, how DNA sites may determine dimerization preferences^53^ and residue-level details of DNA binding and search mechanisms^55,56,100^. Some research publications describe their observations as unusual in the context of the preferred monomer search pathway. For example, most free protein in their solutions being dimeric^50^, or failure to observe monomer-DNA intermediates under equilibrium conditions^57^. The kinetic preference is also referenced in reviews^59–61^ and patents pertaining to peptidomimetics. Our simplified equilibrium and kinetic schemes for the bZIP CREB form a basis for modelling and understanding important features of the search process. In our DNA mixtures both monomer and dimer pathways operate simultaneously, but the dimer pathway is overwhelmingly favoured even when abundant non-target DNA is present. Therefore, our results call into question the existing narrative regarding the active search species of oligomeric transcription factors.

## ACKNOWLEDGEMENTS

UK Medical Research Council [MR/N024168/1 to S.L.S.]; C.K. was supported by the UK BBSRC DTP scheme [DDT00060]. The authors would like to thank Hisayo Sadamoto for personal discussion of their published work, and Nobuhiko Yamamoto for provision of original immunostaining data from their published work.

