## Supplementary Materials for "A basic leucine zipper uses a dimer pathway to locate its targets in DNA mixtures"

**Table S1.** Kinetic and thermodynamic parameters for the dimerization reaction of several CREB bZIP constructs used in this study.

| Construct | $K_{d,M\text{-}M}$ (μM) | $k_{+,\,M\text{-}M}$ ($\mu M^{-1}s^{-1}$) | $k_{-,\,M\text{-}M}$ ($s^{-1}$) |
|---|---|---|---|
| CREB-bZIP | 0.0057 ± 0.0003 | 10 ± 2 | 0.029 ± 0.006 |
| CREB-bZIP-L318V | 14 ± 2 | 3.2 ± 0.4 | 21.4 ± 1.3 |
| CREB-bZIP-L332V | 0.91 ± 0.14 | 0.44 ± 0.04 | 0.201 ± 0.013 |

**Table S2.** Equilibrium dissociation constants ($K_d$) used in equilibrium modelling (Fig. 3), based on estimated values from this study for CREB-bZIP.

| Equilibrium dissociation constant | Value* |
|---|---|
| $K_{d,M\text{-}M}$ | (5.7 ± 0.3) nM |
| $K_{d,M\text{-}CRE}$ | (210 ± 40) μM |
| $K_{d,M\text{-}Comp}$ | (440 ± 30) μM |
| $K_{d,D\text{-}CRE}$ | (214 ± 6) pM |
| $K_{d,D\text{-}Comp}$ | (158 ± 9) nM |

*Values are those obtained for AlexaFluor®488-DNA constructs, however competition equilibrium studies for CREB-bZIP-FD with unlabelled DNA constructs (Fig. S9) indicate differences due to labelling are minor (< 2-fold). Since not all $K_d$ values can be determined without an extrinsic fluorophore labelled values are used throughout for consistency.

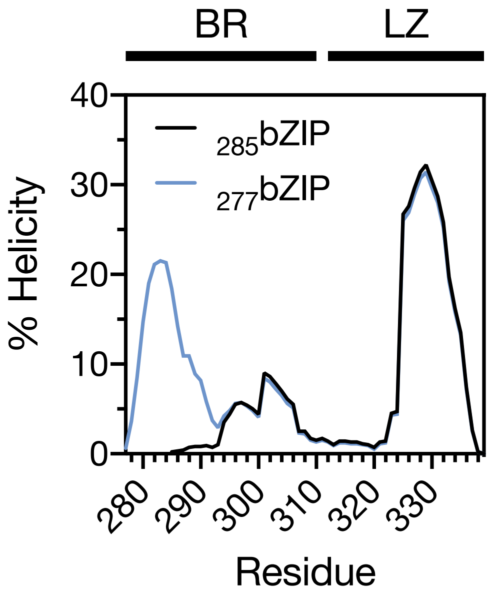


**Fig. S1. AGADIR predictions for CREB-bZIP and an N-terminally truncated version used in previous studies.** Predicted helicity for the N-terminal residues of CREB 285-315 ($_{285}$bZIP, black) is increased with additional N-terminal residues to form CREB 277-315 ($_{277}$bZIP, blue) according to the helical predictor AGADIR[1]. Constructs in this manuscript are based on CREB 277-315, which is referred to as CREB-bZIP throughout.

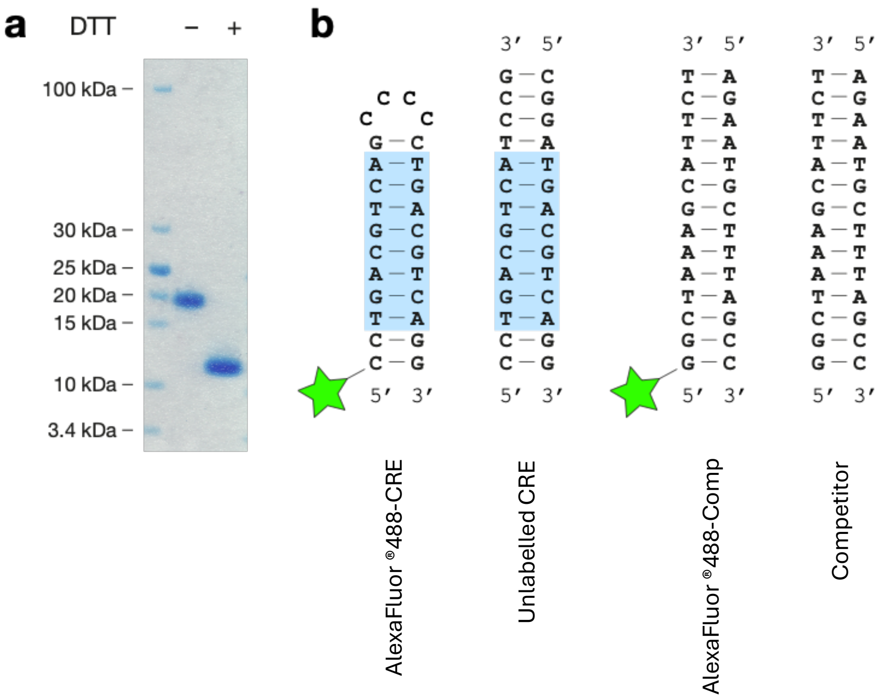


**Fig. S2. Main components of mixing studies. a,** SDS-PAGE analysis of CREB-bZIP-FD. The samples containing 20 µM disulphide-linked protein were mixed with either DTT (final concentration, 10 mM) or an equal volume of water, incubated at room temperature for 30 min and separated on 4-12% polyacrylamide gel. The results confirm homogeneity of protein preparation, as well as the presence of disulphide bond. **b,** DNA oligonucleotides used in the binding experiments. CRE site is highlighted in blue, 5'-AlexaFluor®488 label is drawn as star. Alexa Fluor® 488-CRE, labelled self-annealing hairpin CRE; unlabelled CRE, unlabelled linear double-stranded CRE used in dissociation studies; Alexa Fluor® 488-Comp, labelled competitor DNA used to characterise binding interactions with non-target competitor DNA; competitor, unlabelled competitor DNA used in excess in target search studies.

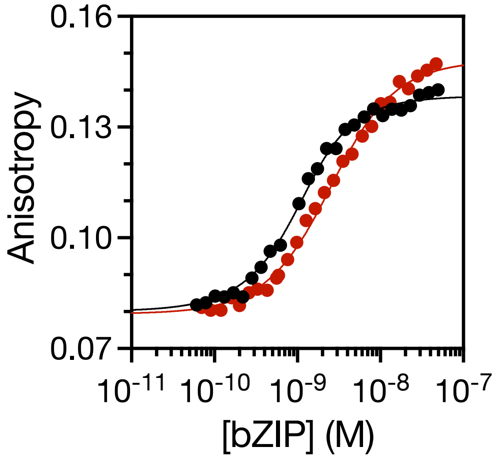


**Fig. S3. Equilibrium binding curves of CREB-bZIP-FD (black) and a weakly binding mutant CREB-bZIP-FD R285_K285insG$_3$ (red) to Alexa Fluor® 488-CRE DNA.** The lines are the fits to Eq. 3. Binding of CREB-bZIP-FD can be described as sub-nanomolar but cannot be estimated accurately since the probe concentration (1 nM AlexaFluor®488-CRE DNA) is larger than $K_{d,D\text{-}CRE}$.

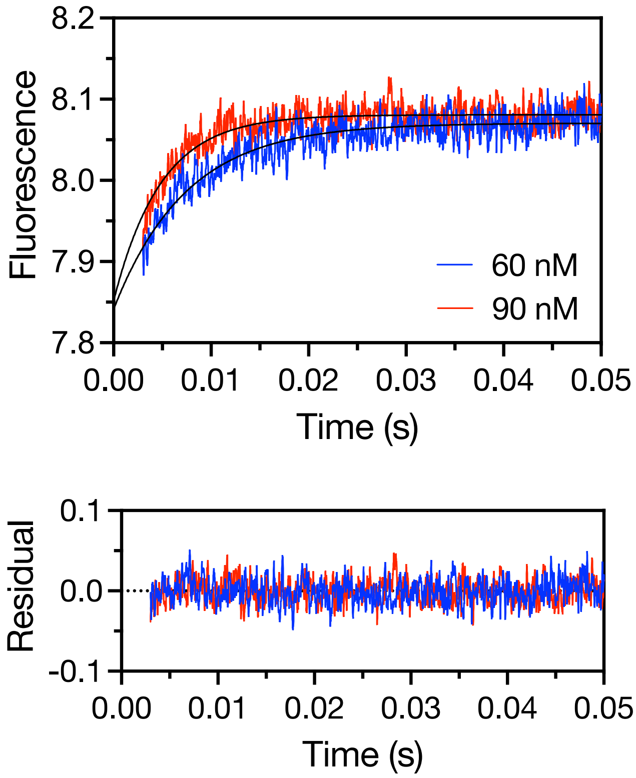


**Fig. S4. Representative stopped-flow kinetic traces for association of CREB-bZIP-FD protein with 10 nM AlexaFluor®488-CRE DNA.** The concentration of CREB dimer in the association experiment was 60 and 90 nM. Solid lines are fits to a single-exponential decay function.

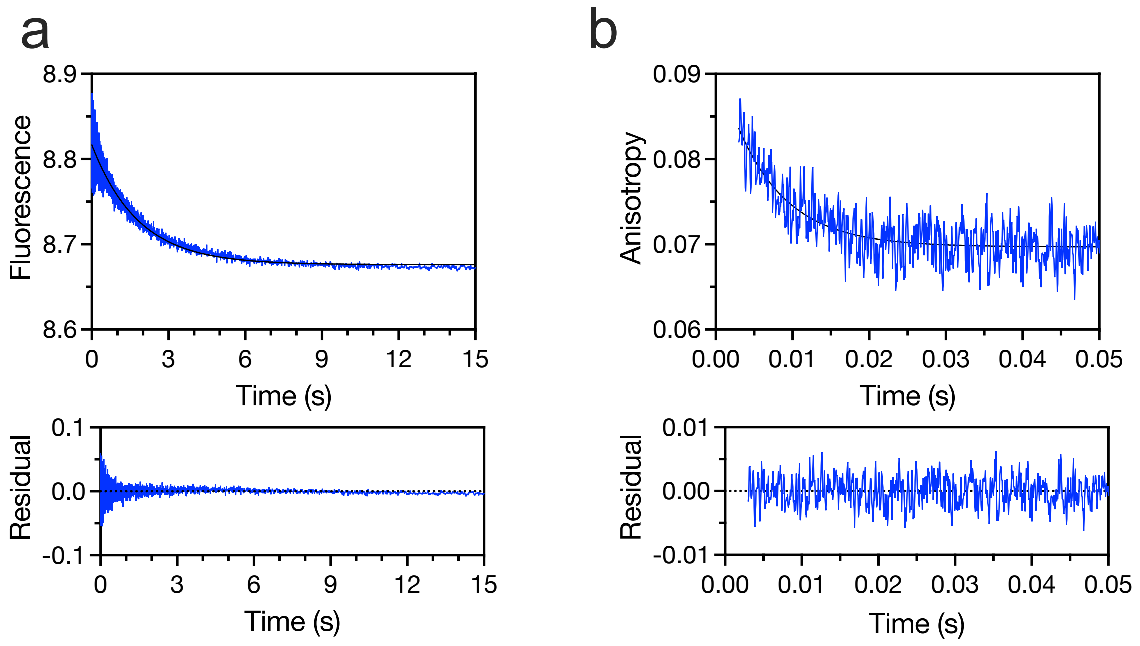


**Fig. S5. Representative stopped-flow kinetic traces for irreversible dissociation** of **a.** 100 nM CREB-bZIP-FD and 5nM AlexaFluor®488-CRE **b.** 200 nM CREB-bZIP-FD and 200 nM AlexaFluor®488-Comp. In **b** fluorescence anisotropy was used since no change in intensity was observed upon binding.

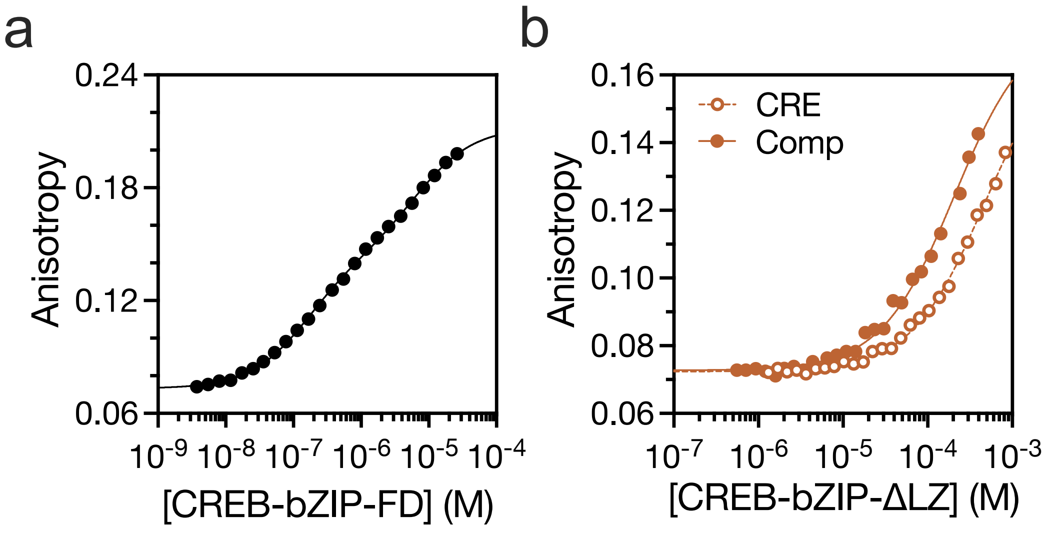


**Fig. S6.** Fluorescence anisotropy equilibrium binding curves of CREB-bZIP-FD (**a**) and CREB-bZIP-ΔLZ to AlexaFluor®488-CRE (open circles) and AlexaFluor®-488-Comp (closed circles) (**b**). DNA concentration was 5 nM. The lines are the fits to Eq.4 and Eq. 3 in Methods.

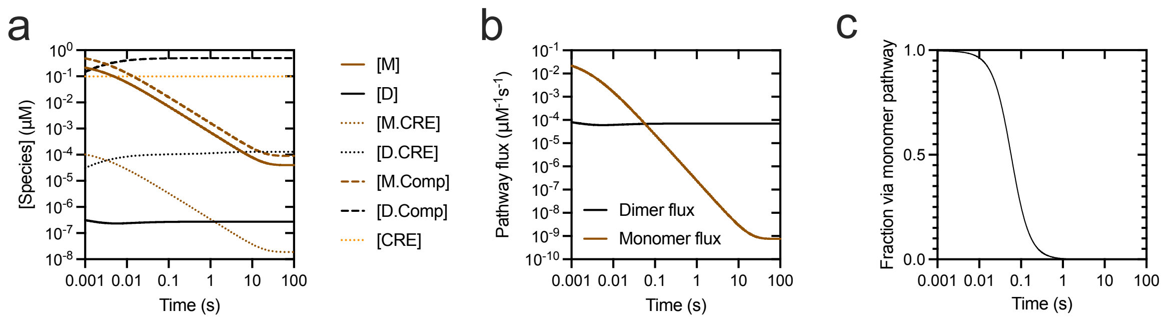


**Fig. S7. Kinetic modelling of species concentrations in mixtures of CREB and DNA demonstrates preference for dimer pathway is established quickly even for maximally effective traps.** Initial conditions are deliberately chosen to favour the monomer pathway with all CREB being monomeric. The rate constant for dimer dissociation from competitor is matched to that for CRE to generate a maximum possible kinetic trap effect ($k_{-,\mathrm{D\text{-}Comp}} = k_{-,\mathrm{D\text{-}CRE}}$) but otherwise concentrations and rate constants match those in Fig. 4a-c: [CREB] = 1 μM, [CRE] = 0.1 μM, [Comp] =1000 μM (chosen to approximate those found within the eukaryotic cell nucleus), $k_{+,\mathrm{M\text{-}M}}$ = 10 $\mu M^{-1}s^{-1}$, $k_{-,\mathrm{M\text{-}M}}$ = 0.029 $s^{-1}$, $k_{+,\mathrm{D\text{-}CRE}}$ = 2570 $\mu M^{-1}s^{-1}$, $k_{-,\mathrm{D\text{-}CRE}}$ = 0.55 $s^{-1}$, $k_{+,\mathrm{D\text{-}Comp}}$ = 1000 $\mu M^{-1}s^{-1}$, $k_{+,\mathrm{M\text{-}CRE}} = k_{+,\mathrm{M\text{-}Comp}} = k_{+,\mathrm{M\text{-}M.CRE}} = k_{+,\mathrm{M\text{-}M.Comp}}$ = 1000 $\mu M^{-1}s^{-1}$, $k_{-,\mathrm{M\text{-}CRE}}$ = 210 μM $k_{+,\mathrm{M\text{-}CRE}}$, $k_{-,\mathrm{M\text{-}Comp}}$ = 440 μM $k_{+,\mathrm{M\text{-}Comp}}$. Values for $k_{-,\mathrm{M\text{-}M.CRE}}$ and $k_{-,\mathrm{M\text{-}M.Comp}}$ were set as required to maintain thermodynamic cycles. Evolution of species concentration (**a**), pathway flux (**b**) and fractional flux through the monomer pathway (**c**) for these values of variables demonstrate that the dimer pathway quickly overtakes the monomer pathway in terms of flux. Note that this scenario is for illustrative purposes only and is highly unrealistic: the lack of specificity implied for dimeric CREB is seen to result in minimal occupation of the CRE bound state.

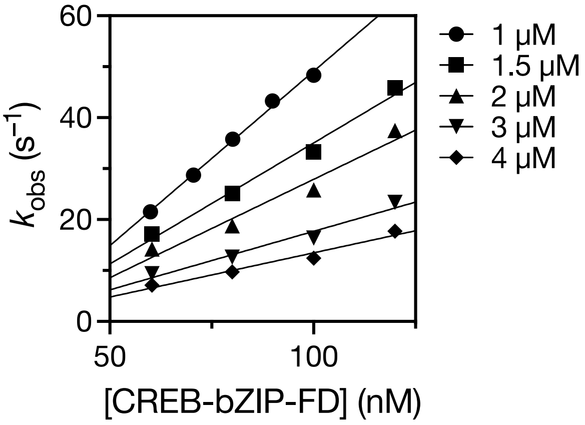


**Fig. S8. Retardation in target search due to presence of (non-target) competitor DNA.** Observed rate constants for binding of excess CREB-bZIP-FD with 10 nM AlexaFluor488®-CRE in the presence of different concentrations of unlabelled competitor remain linearly dependent on protein concentration. The apparent rate constant was determined as the gradient of these lines and decreases with competitor concentration. Protein concentration is that for dimer.

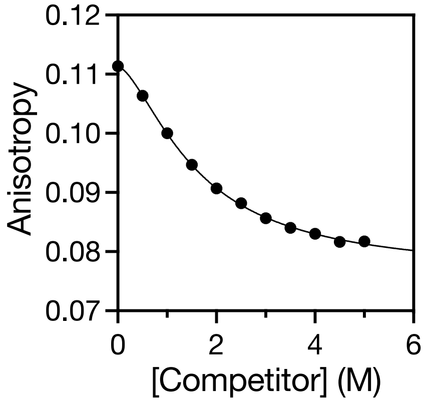


**Fig. S9. Competition equilibrium curve to estimate binding affinity of CREB-ZIP-FD with unlabelled competitor DNA.** Fluorescence anisotropy of pre-formed complexes of CREB-ZIP-FD and Alexa Fluor® 488-Comp DNA upon incubation with various concentrations of unlabelled competitor DNA (Fig. S2b). Concentrations of CREB-bZIP-FD and AlexaFluor®488-Comp DNA were 500 nM and 100 nM respectively. This competition equilibrium curve was fit to Eq. 6 to extract the binding affinity to unlabelled competitor DNA as (300 ± 30) μM.

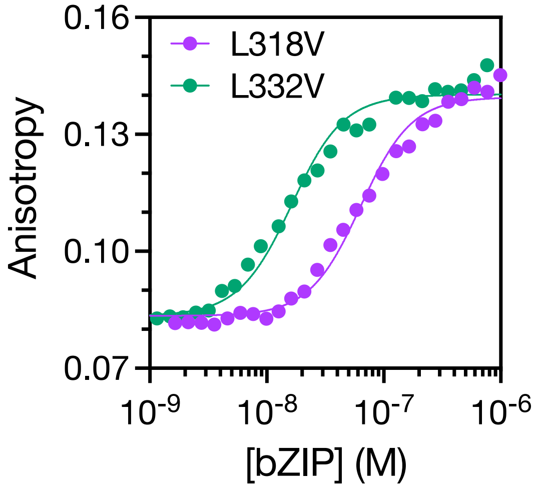


**Fig. S10. Zipper destabilisation significantly decreases target DNA binding affinity to extent expected for a dimer-only binding model.** Equilibrium binding of CREB-bZIP-L318V and CREB-bZIP-L332V to 1 nM AlexaFluor®488-CRE DNA. The data was fit to Eq. 5, suitable for a reversible reaction scheme where protein dimerises and then binds to DNA (a dimer-only binding model); $K_{D,\,D\text{-}CRE}$ was constrained to the value obtained for CREB-bZIP-FD (214 pM) to obtain $K_{d,M\text{-}M}$ of (16 ± 2) µM for L318V and (0.91 ± 0.11) µM for L332V. These are the same within error as those determined in the dilution experiment of (14 ± 2) µM (L318V) and (0.91 ± 0.14) µM (L332V).

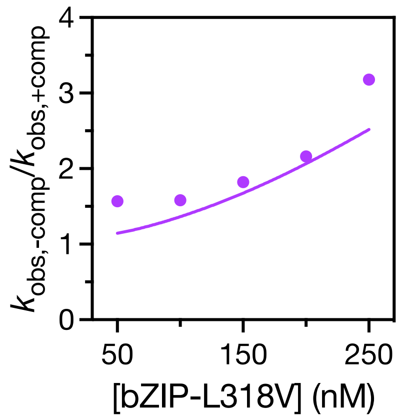


**Fig. S11. Predicted retardation of association rate constant on introduction of competitor is consistent with dimer-only binding pathway.** Ratio of observed rate constants for association of CREB-bZIP-L318V with 5 nM AlexaFluor®488-CRE from fluorescence stopped-flow spectroscopy performed in the absence and presence of 2 µM competitor DNA (purple circles). The ratio is observed to decrease with protein concentration. The solid line represents the prediction based on numerical integration of the rate equations for a dimer-only binding model and independently obtained estimates for the six rate constants.

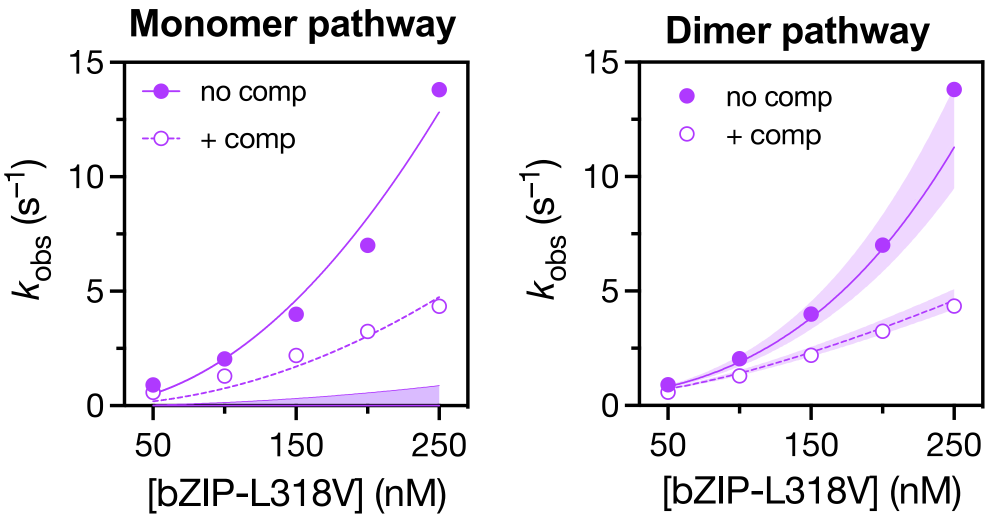


**Fig. S12. Observed rate constants for association of CREB-bZIP-L318V with 5 nM AlexaFluor®488-CRE from fluorescence stopped-flow spectroscopy better match the predictions of a dimer-only binding model.** Experiments were performed in the absence (open circles) and presence (closed circles) of 2 µM competitor DNA. The predicted range of rate constants for monomer and dimer pathways are shown as purple shading in the left and right panels respectively. Observed rate constants are significantly higher than those predicted by the monomer pathway for a $k_{+,\mathrm{M\text{-}M.CRE}}$ of 3 $\mathrm{nM^{-1}s^{-1}}$. In the left panel both datasets are fit to a quadratic equation. In the absence of competitor the monomer-only binding model predicts $k_{obs} = \frac{k_{+,M-M.CRE}}{K_{d,M-CRE}}[bZIP]^2 + k_{-,M-M.CRE}$, which would imply an extremely high $k_{+,\mathrm{M\text{-}M.CRE}}$ of 30-60 $\mathrm{nM^{-1}s^{-1}}$. In the right panel the solid lines are the predicted rate constants (not fitted) for a dimer-only binding model based on numerical integration of the rate equations for a dimer-only binding model and independently obtained estimates for the six rate constants.

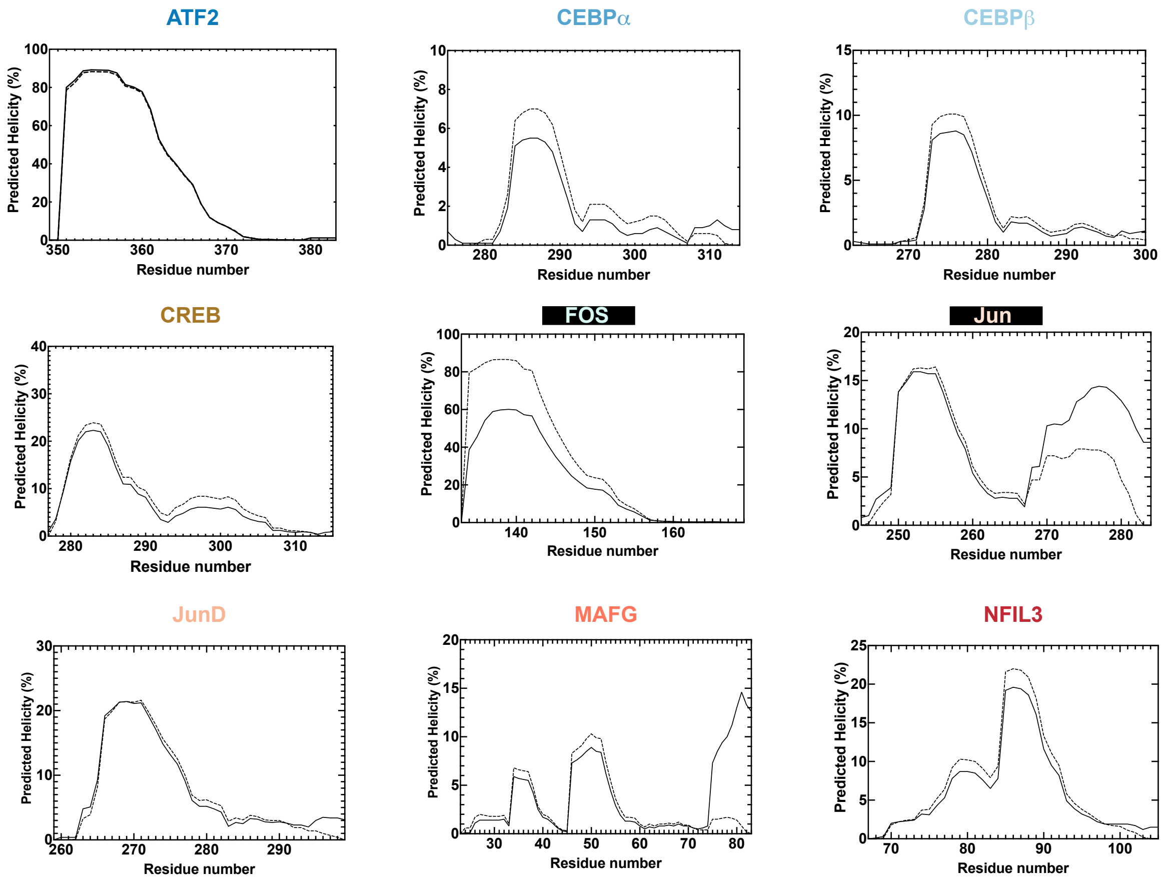


**Fig. S13. AGADIR predictions for all forced monomeric (bZIP-ΔLZ) peptides.** Solid lines are predictions for full-length protein, whereas dotted lines are for the truncated bZIP-ΔLZ used in the study. The monomeric peptides all have very similar, or slightly higher, levels of predicted helicity than the full-length protein.

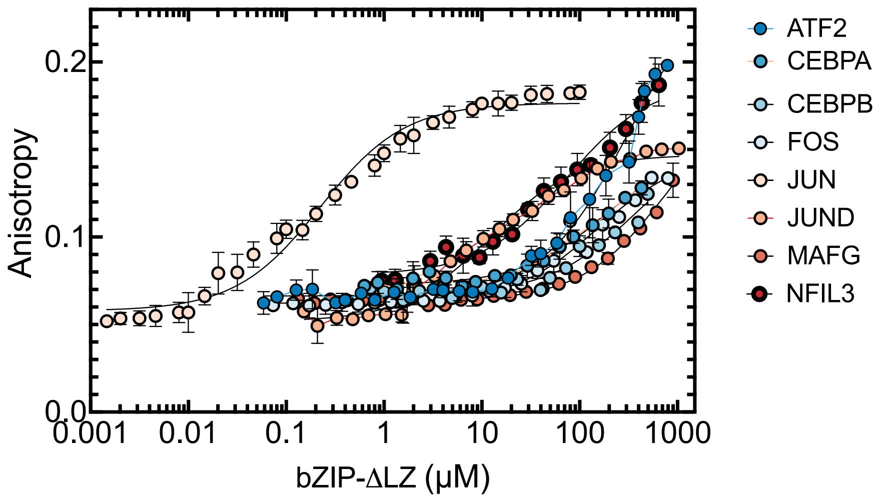


**Fig. S14. Most monomeric bZIP peptides have poor target binding affinity.** Fluorescence anisotropy equilibrium curve for binding of monomeric bZIP-ΔLZ peptides lacking the dimerization domain to 5 nM 5'-labelled AlexaFluor®488 target-containing DNA. DNA targets are as outlined in Materials and Methods. Solid black lines represent the best fit to Eq. 3.

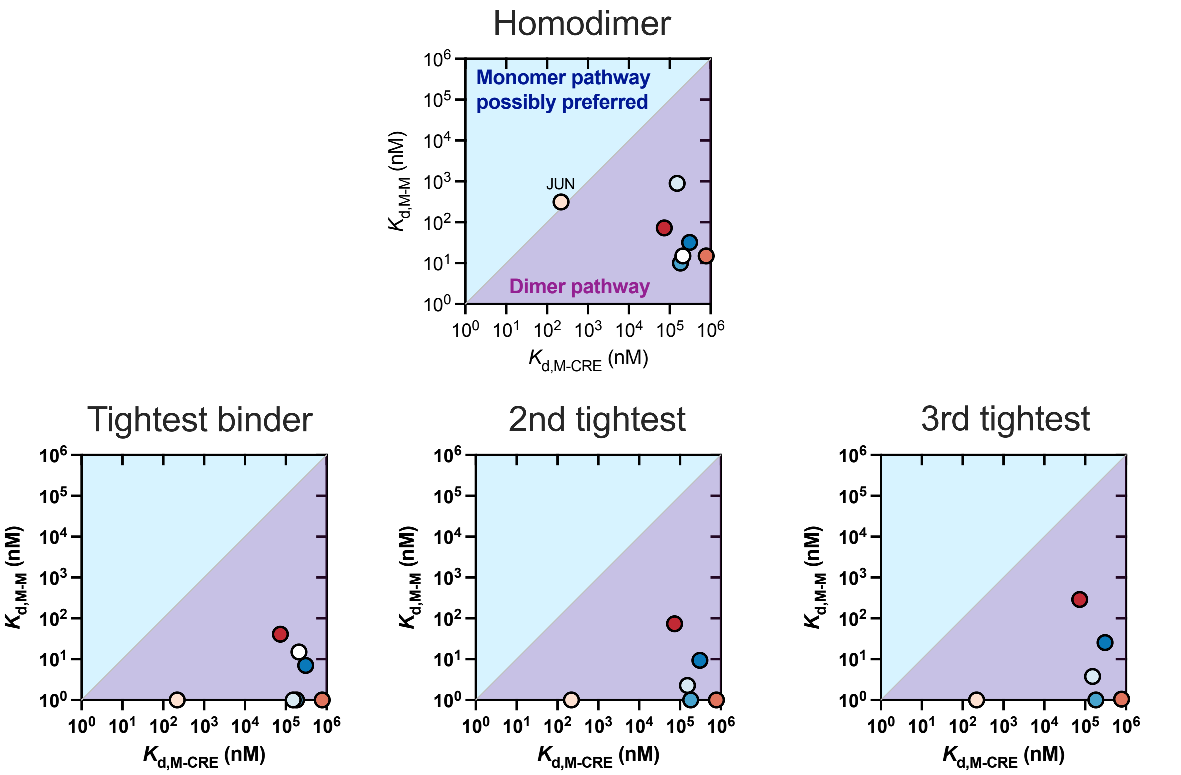


**Fig. S15. Comparison of the monomer binding affinity for target DNA obtained in this study with those for dimerization estimated previously by Reinke et al. by microarray methods[2].** At equilibrium the proportion of target complex formed through the monomer pathway is $\varphi_M \sim \frac{1}{\frac{K_{d,M-Tar}}{K_{d,M-M}}+1}$ (Materials and Methods). Purple shading represents the case when $K_{d,M\text{-}M} > K_{d,M\text{-}Tar}$ and blue shading represents the case when $K_{d,M\text{-}M} < K_{d,M\text{-}Tar}$. Except for Jun, $K_{d,M\text{-}M}$ are orders of magnitude smaller than $K_{d,M\text{-}Tar}$ so relatively little flux is expected through the monomer pathway at equilibrium. bZIP proteins form homodimers and heterodimers as part of a complex network, though CREB is described to form exclusive homodimers[2]. The upper panel shows equilibrium dissociation constant for bZIP monomer binding itself ($K_{d,M\text{-}M}$), and lower panels show those for binding to its three most favoured binding partners to form heterodimers ($K_{d,M\text{-}M*}$).

### EQUATION DERIVATIONS

#### Derivation of equation for equilibrium dissociation of protein dimer

Let *M* denote the protein monomer and *D* – the protein dimer. Dissociation of protein can be described by the following chemical equation:

$$2M \leftrightharpoons D. \qquad \text{(Eq. S1)}$$

Let [*M*] and [*D*] denote the concentrations of the respective species, and $P_T$ denote the total concentrations of protein. The mass balance gives

$$P_T = [M] + 2[D]. \qquad \text{(Eq. S2)}$$

Let $K_{d,M\text{-}M}$ denote the dissociation constant of the dimer. When the system reaches equilibrium,

$$K_{d,M-M} = \frac{[M]^2}{[D]}. \qquad \text{(Eq. S3)}$$

Combining the above two equations and simplifying the result gives:

$$2[M]^2 + K_{d,M-M}[M] - K_{d,M-M}P_T = 0. \qquad \text{(Eq. S4)}$$

The value of [*M*] is a positive root of Eq. S4:

$$[M] = \frac{-K_{d,M-M} + \sqrt{K_{d,M-M}{}^2 + 8K_{d,M-M}P_T}}{4} = K_{d,M-M}\left(\sqrt{\frac{1}{16} + \frac{1}{2}\frac{P_T}{K_{d,M-M}}} - \frac{1}{4}\right). \qquad \text{(Eq. S5)}$$

The overall fluorescence intensity of a mixture *I* is the sum of individual intensities of emitting species $I_i$ multiplied by their molar concentrations $c_i$:

$$I = \sum I_i c_i = I_m[M] + I_d[D]. \qquad \text{(Eq. S6)}$$

Eqs. S2 and S6 can be combined and rearranged as follows:

$$I^* = I_d^* + \Delta I^* \left(\frac{[M]}{P_T}\right), \qquad \text{(Eq. S7)}$$

where

$$I^* = \frac{I}{P_T},$$ (Eq. S7a)

$$I_d^* = \frac{1}{2} I_d,$$ (Eq. S7b)

$$\Delta I^* = I_m - I_d^*.$$ (Eq. S7c)

Now, we can substitute Eq. S5 into Eq. S7 to obtain the final formula expressing the fluorescence intensity through the $K_{d,M-M}$ and total concentration of protein:

$$I^* = I_d^* + \Delta I^* \left( \frac{K_{d,M-M}}{P_T} \left( \sqrt{\frac{1}{16} + \frac{1}{2} \frac{P_T}{K_{d,M-M}}} - \frac{1}{4} \right) \right).$$ (Eq. S8)

**Deviation of equation for dissociation rate constant of protein dimer**

Let $k_1$ and $k_2$ denote the association and dissociation rate constants, respectively, for homodimerization reaction Eq. S1. The rate of change in monomer concentration can be expressed as:

$$\frac{d[M]}{dt} = -k_{+,M=M}[M]^2 + 2k_{-,M-M}[D]$$

(Eq. S9)

Identifying the equilibrium constant $K_{d,M-M}$ as

$$K_{d,M-M} = 2\frac{k_{-,M-M}}{k_{+,M-M}}$$

(Eq. S10)

and requiring mass balance (Eq. S2) leads to:

$$-k_{+,M-M}dt = \frac{d[M]}{[M]^2 + \frac{K_{d,M-M}}{2}[M] - \frac{K_{d,M-M}}{2}P_T}$$

(Eq. S11)

Finding the roots of the quadratic expression in the denominator allows this to be rewritten as:

$$-k_{+,M-M}dt = \frac{d[M]}{\left([M]+\frac{K_{d,M-M}+2z}{4}\right)\left([M]+\frac{K_{d,M-M}-2z}{4}\right)},$$ (Eq. S12)

where

$$z = \sqrt{\frac{K_{d,M-M}^{2}}{4} + 2K_{d,M-M}P_T} = 2K_{d,M-M}\sqrt{\frac{1}{16} + \frac{1}{2}\frac{P_T}{K_{d,M-M}}}.$$ (Eq. S12a)

Eq. S12 may then be integrated to obtain an expression for the concentration of monomeric protein with time:

$$[M] = \frac{\frac{K_{d,M-M}-2z}{4} - \left(\frac{K_{d,M-M}+2z}{4}\right)Ne^{-k_{+,M-M}zt}}{Ne^{-k_{+,M-M}zt}-1},$$ (Eq. S13)

where $N$ is a constant of integration that may be written in terms of the initial monomer concentration $[M]_0$ by evaluation of Eq. S13 at time zero:

$$N = \frac{\frac{K_{d,M-M}-2z}{4}+[M]_0}{\frac{K_{d,M-M}+2z}{4}+[M]_0}.$$ (Eq. S14)

$[M]_0$ is related to equilibrium concentration of monomer $[M]_{\text{syringe}}$ in syringe prior to 11-fold dilution defined by Eq. S5, hence:

$$[M]_0 = \frac{[M]_{\text{syringe}}}{11} = \frac{K_{d,M-M}}{11}\left(\sqrt{\frac{1}{16} + \frac{11}{2}\frac{P_T}{K_{d,M-M}}} - \frac{1}{4}\right).$$ (Eq. S15)

By considering limits, one can identify:

$$[M]_\infty = -\left(\frac{K_{d,M-M}-2z}{4}\right) = K_{d,M-M}\left(\sqrt{\frac{1}{16} + \frac{1}{2}\frac{P_T}{K_{d,M-M}}} - \frac{1}{4}\right),$$ (Eq. S16)

where $[M]_\infty$ denotes the monomer concentration at equilibrium. Substituting of Eq. S16 into Eq. S13 yields:

$$[M] = \frac{[M]_\infty - ([M]_\infty - z)Ne^{-k_{+,M-M}zt}}{1 - Ne^{-k_{+,M-M}zt}} = [M]_\infty + z\left(\frac{e^{-k_{+,M-M}zt}}{\frac{1}{N} - e^{-k_{+,M-M}zt}}\right).$$ (Eq. S17)

Finally, Eq. S10 allows to express $-k_{+,M-M}z$ in Eq. S17 through $k_{-,M-M}$:

$$k_{+,M-M}z = k_{-,M-M}\sqrt{1+\frac{8P_T}{K_{d,M-M}}}. \quad \text{(Eq. S18)}$$

**Deviation of equation for equilibrium binding of a pre-formed protein dimer to DNA**

Let *CRE* denote the DNA molecule and *D.CRE* – the protein-DNA complex. The binding of protein to DNA in our system can be described by the following set of equations:

$$2M \leftrightharpoons D, \quad \text{(Eq. S19)}$$

$$CRE + D \leftrightharpoons D.CRE. \quad \text{(Eq. S20)}$$

Let $[CRE]$ and $[D.CRE]$ denote the concentrations of the respective species, and $CRE_T$ denote the total concentrations of protein and DNA. The mass balance gives

$$P_T = [M] + 2[D] + 2[D.CRE], \quad \text{(Eq. S21)}$$

$$CRE_T = [CRE] + [D.CRE]. \quad \text{(Eq. S22)}$$

Let $K_{d,D\text{-}CRE}$ denote dissociation constants of the protein-DNA complex and $K_{d,M\text{-}M}$ denote dissociation constant of the protein dimer. When the system reaches equilibrium,

$$K_{d,M-M} = \frac{[M]^2}{[D]}, \quad \text{(Eq. S23)}$$

$$K_{d,D-CRE} = \frac{[D][CRE]}{[D.CRE]}. \quad \text{(Eq. S24)}$$

Substitution of Eqs. S22-S24 into Eq. S21 yields an expression for $[D.CRE]$:

$$[D.CRE] = \frac{1}{2}P_T - \frac{1}{2}\sqrt{K_{d,M-M}\frac{K_{d,D-CRE}[D.CRE]}{CRE_T-[D.CRE]}} - \frac{K_{d,D-CRE}[D.CRE]}{CRE_T-[D.CRE]}. \quad \text{(Eq. S25)}$$

The anisotropy of a mixture *r* is the sum of individual anisotropies of emitting species $r_i$ weighted by their fractional intensities $f_i$ as described by Lakowicz[6]:

$$r = \sum r_i f_i = r_{CRE} f_{CRE} + r_{D.CRE} f_{D.CRE}, \quad \text{(Eq. S26)}$$

where

$$f_{CRE} + f_{D.CRE} = 1. \quad \text{(Eq. S27)}$$

Eqs. S26 and S27 can be combined and rearranged as follows:

$$r = r_{CRE} + \Delta r f_{D.CRE}, \qquad \text{(Eq. S28)}$$

where

$$\Delta r = r_{D.CRE} - r_{CRE}. \qquad \text{(Eq. S28a)}$$

If binding of protein to DNA does not affect its fluorescence intensity, the fractional intensities can be re-written as molar fractions:

$$r = r + \frac{\Delta r[D.CRE]}{CRE_T}. \qquad \text{(Eq. S29)}$$

Otherwise, the anisotropy can be adjusted as described by Crabtree and Shammas[7]. Substituting Eq. S25 into Eq. S29 gives

$$r = r_{CRE} + \frac{\Delta r}{CRE_T}\left(\frac{1}{2}P_T - \frac{1}{2}\sqrt{K_{d,M-M}\frac{K_{d,D-CRE}[D.CRE]}{CRE_T - [D.CRE]}} - \frac{K_{d,D-CRE}[D.CRE]}{CRE_T - [D.CRE]}\right). \qquad \text{(Eq. S30)}$$

Solving Eq. 29 for $[D.CRE]$ and substituting into Eq. S30 yields the final formula:

$$r = r_{CRE} + \frac{\Delta r}{CRE_T}\left(\frac{1}{2}P_T - \frac{1}{2}\sqrt{K_{d,M-M}\frac{K_{d,D-CRE}\,y}{CRE_T - y}} - \frac{K_{d,D-CRE}\,y}{CRE_T - y}\right), \qquad \text{(Eq. S31)}$$

where

$$y = \frac{CRE_T(r - r_{CRE})}{\Delta r}. \qquad \text{(Eq. S31a)}$$

It's worth noting that Eq. S31 is defined in an implicit form as $r = f(P_T, r)$ so the solution for $r$ was obtained using numerical methods.
